# Cytokines reprogram airway sensory neurons in asthma

**DOI:** 10.1101/2023.01.26.525731

**Authors:** Théo Crosson, Shreyas Bhat, Jo-Chiao Wang, Clara Salaun, Eleanne Fontaine, Katiane Roversi, Herbert Herzog, Moutih Rafei, Rikard Blunck, Sebastien Talbot

**Affiliations:** Département de Pharmacologie et Physiologie, Université de Montréal, Canada; Centre Interdisciplinaire sur le Cerveau et l’Apprentissage, Université de Montréal, Canada; Département de Physique, Université de Montréal, Canada; Gavan Institute of Medical Research, Australia; Department of Physiology and Pharmacology, Karolinska Institutet. Sweden.; Department of Biomedical and Molecular Sciences, Queen’s University. Canada.

**Keywords:** Allergy, asthma, IL-6, IL-13, neuro-immunology, nociceptor neurons, NPY1R, vagus nerve

## Abstract

Nociceptor neurons play a crucial role in maintaining the body’s homeostasis by detecting and responding to potential dangers in the environment. However, this function can be detrimental during allergic reactions, since vagal nociceptors can contribute to immune cell infiltration, bronchial hypersensitivity, and mucus imbalance, in addition to causing pain and coughing. Despite this, the specific mechanisms by which nociceptors acquire pro-inflammatory characteristics during allergic reactions are not yet fully understood. In this study, we aimed to investigate the molecular profile of airway nociceptor neurons during allergic airway inflammation and identify the signals driving such reprogramming. Using retrograde tracing and lineage reporting, we identified a unique class of inflammatory vagal nociceptor neurons that exclusively innervate the airways. In the ovalbumin mouse model of airway inflammation, these neurons undergo significant reprogramming characterized by the upregulation of the NPY receptor *Npy1r*. A screening of cytokines and neurotrophins revealed that IL-1β, IL-13 and BDNF drive part of this reprogramming. IL-13 triggered *Npy1r* overexpression in nociceptors via the JAK/STAT6 pathway. In parallel, sympathetic neurons and macrophages release NPY in the bronchoalveolar fluid of asthmatic mice, which limits the excitability of nociceptor neurons. Single-cell RNA sequencing of lung immune cells has revealed that a cell-specific knockout of Npy1r in nociceptor neurons in asthmatic mice leads to an increase in airway inflammation mediated by T cells. Opposite findings were observed in asthmatic mice in which nociceptor neurons were chemically ablated. In summary, allergic airway inflammation reprograms airway nociceptor neurons to acquire a pro-inflammatory phenotype, while a compensatory mechanism involving NPY1R limits nociceptor neurons’ activity.

## BACKGROUND

Sensory neurons, also known as nociceptor neurons, are heterogenous and can be differentiated based on their expression profiles ^1–3^, degree of myelination ^4^, the type of cues to which they are sensitive ^4,5^, the reflexes that they initiate, their anatomical location ^6^ or the organ that they innervate ^7,8^. For instance, internal organs such as the lungs are innervated by the vagus nerve^9–12^, whose sensory neurons originate from the jugular-nodose complex ganglia (JNC).

Sensory neurons form a key line of defense against environmental dangers. They detect a broad range of thermal, mechanical, and chemical threats and respond by means of protective reflexes such as the itch and cough reflexes^13,14^. To fulfill their role in host defense, nociceptors are geared to detect and respond to a variety of danger signals ranging from cytokines to inflammatory lipids, allergens, fungi, cancer cells, bacteria, toxins, and even immunoglobulins^5,15,16^. Via the axon reflex, nociceptors respond by locally releasing neuropeptides. While these peptides can have direct inflammatory and immunomodulatory actions^17–23^, nociceptors’ impact on inflammation is also indirect, via modulation of the autonomic nervous system^24^.

Our work, and the one of others, has previously shown that nociceptor neurons regulate airway^25,26^, skin^14,27^ and eye^28,29^ allergic inflammation in models ranging from ovalbumin^30–33^, HDM^18,19,34^, candidiasis^21^ vitamin D^35,36^, poison ivy/IL-33^37^, as well as proteases^20,38^. In the context of asthma, vagal nociceptors neurons worsen airway hyperreactivity by promoting bronchoconstriction, coughing, mucus imbalance, and immune cell infiltration^18,30,39,40^. Likewise, numerous studies have indicated that neurons can detect a variety of cytokines^14,16,18,37,41–43^. Furthermore, it has been demonstrated that cytokine-induced intracellular signaling in sensory neurons involves JAK1^14,27^, although this area requires further research.

Given that IL-4 and IL-13 are crucial cytokines released during Th2 inflammation, including in response to house dust mite (HDM) challenges, we posit that similar reprogramming of airway neurons likely happens in this context. Here we sought to define the molecular profile of airway nociceptor neurons during allergic airway inflammation and to identify the signals driving their reprogramming. Using retrograde tracing and lineage reporting, we first identified the populations of nociceptor neurons that innervates the airways. Then, upon allergic airway inflammation, we found a drastic reprogramming of airway nociceptor neurons characterized by the upregulation of the NPY receptor *Npy1r.* Next, we identified that the asthma-driving cytokine IL-13, via JAK/pSTAT6 signaling, drives part of this transcriptional reprogramming. Along with *Npy1r* overexpression, we found that NPY was elevated in the bronchoalveolar fluid of asthmatic mice and expressed by sympathetic neurons and M2 macrophages. In cultured JNC neurons, NPY1R agonist decreased nociceptor neurons cAMP and blunted neuronal excitability. Finally, knocking out *Npy1r* in nociceptors altered T cells gene expression in the lung, in an opposite fashion to the one observed with nociceptor neuron chemoablation.

## RESULTS

### A class of inflammatory nociceptor specifically innervates the airways

Via vagal projections, the JNC innervates most visceral organs^44^. Single-cell RNA sequencing datasets revealed that JNC neurons are highly heterogenous^2,3,10^ and provide a transcriptome-based neuron subtype classification^2,3,6,45^. While genetically guided optogenetic studies identified JNC neuronal subtypes controlling breathing and tracheal reflexes^1,10,46^, additional molecular characterization of airway nociceptors in inflammatory context is lacking.

To address this, we tracked airway nociceptors using the Na_V_1.8^cre^::tdTomato^fl/wt^ nociceptor neuron reporter mice, in which, we back-labeled airway neurons using the retrograde tracer DiD’ (i.n. 200 µM). Two weeks after DiD’ injection, the JNC neurons were isolated, and airway nociceptor neurons (Na_V_1.8^+^DiD^+^), visceral nociceptor neurons (Na_V_1.8^+^DiD^-^) and a mixed population of Na_V_1.8^-^ cells (e.g. satellite glial cells, macrophages, Nav1.8^neg^ neurons, and stromal cells), were purified by flow cytometry (**Fig. 1A**, **SF. 1A**) and RNA-sequenced (**Fig. 1A-B, SF. 1B**). About 5–10% of JNC nociceptor neurons were labeled with DiD’ (**SF. 1A, C**). Principal component analysis (PCA) analysis showed that the three cell populations were well segregated (**SF. 1B**). This notably implies that airway nociceptors are transcriptionally distinct from other visceral nociceptors. DESeq2 analysis confirmed differential expression of a large number of genes between these two populations (**Fig. 1B-C)**, with the gene coding for the potassium channel *Kcng1* being the most specific marker of airway nociceptors **(Fig. 1B-D)**.

**Figure 1.**
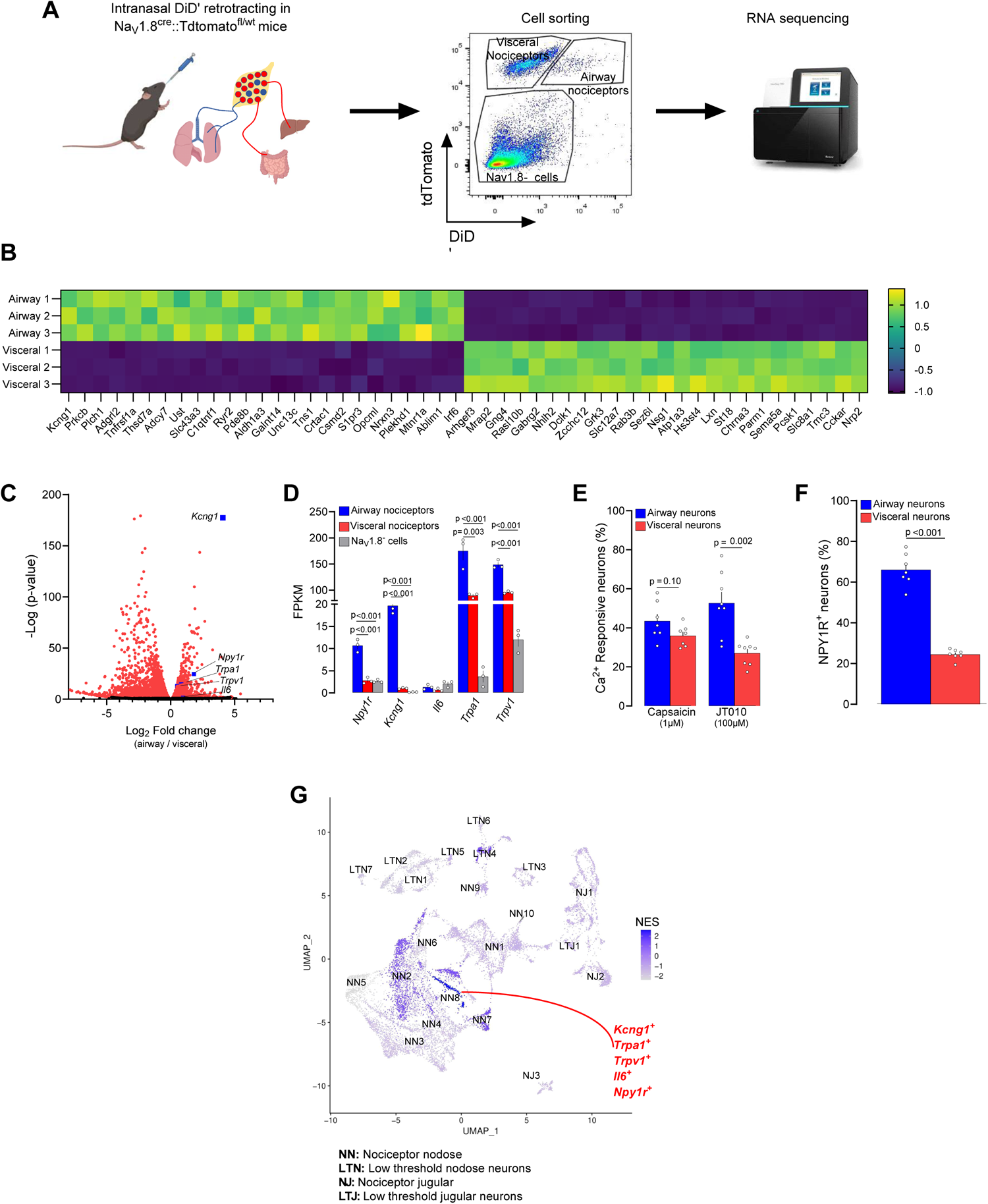
Airway vagal nociceptor neurons have a unique molecular profile. **(A)** Diagram depicting the retrotracing of the airway-innervating nociceptor neurons. To identify airway-innervating nociceptor neurons, naive 8-week-old male and female nociceptor neurons reporter (Na_V_1.8^cre^::tdTomato^fl/wt^) mice were injected intranasally with the retrograde tracer DiD’ (200 µM). Fourteen days later, the mice were euthanized and their JNC ganglia isolated and dissociated. Airway-innervating nociceptor neurons (Na_V_1.8^+^DiD^+^), visceral nociceptors (Na_V_1.8^+^DiD^-^) and Na_V_1.8^-^ cells were purified by flow cytometry and their RNA sequenced (**A**). **(B)** Heatmap displaying the z-score of the 50 top differentially expressed genes between visceral and airway nociceptors. **(C-D)**. A volcano plot of pairwise comparison of airway-innervating nociceptor neurons versus visceral nociceptor neurons shows differentially expressed transcripts in red (adjusted p-value <0.2). Among others, *Npy1r, Kcng1, Trpa1, Trpv1* and *Il6* were enriched in airway-innervating nociceptor neurons (**B**). *Npy1r*, *Kcng1*, *Trpa1* and *Trpv1* were also more expressed in airway nociceptors when compared to Na_V_1.8^-^ cells (**C**). **(E)** 8-week-old male and female C57BL6 mice were injected intranasally with the retrograde tracer DiD’ (200µM). Fourteen days later, the mice were euthanized and their JNC ganglia isolated and dissociated. JNC neurons were cultured (16h) and responsiveness to noxious stimuli was assessed. While responsiveness to the TRPV1 agonist capsaicin (300 nM) was stable between the two groups, the proportion of neurons responsive to the TRPA1 agonist JT010 (50 µM) was higher in airway-innervating neurons (DiD^+^; **D**). **(F)** Naive 8-week-old male and female NPY1R reporter (NPY1R^cre^::tdTomato^fl/wt^) mice were injected intranasally with the retrograde tracer DID’ (200 µM). Fourteen days later, the mice were euthanized and their JNC ganglia isolated and dissociated. JNC neurons were cultured (16h), and neurons defined by KCl calcium responses. Using fluorescent imaging, we found that NPY1R-expressing neurons (tdTomato^+^) are more frequent in the airway-innervating population (DiD^+^; **E**). **(G)** UMAP of *Slc17a6*^+^ (VGLUT2) JNC neurons from single-cell RNA sequencing revealed heterogeneous neuronal subsets. Gene set enrichment analysis was performed to address which neuron subtype preferentially innervated the airways, with normalized enrichment score (NES) indicated in blue. The neuronal cluster NN8 expresses several neuro-inflammatory markers (*Il6, Kcng1, Npy1r, Trpa1, Trpv1*) and exclusively innervates the airways (NES=2.3). Experimental details were defined in *Prescott et al.* and the bioinformatic analysis is described in the method section (**F**). *Data are shown as a heatmap displaying expression z-score (**B**), representative volcano plot displaying DESeq2 normalized count fold change and nominal p-values for each gene **(C)**, as mean ± S.E.M (**D-F**), or as normalized enrichment score (**F**). N are as follows: **B-D**: n=3 biological replicates (4 mice per sample), **E-F**: n=7–8 dishes per group. P-values were determined by DESeq2 analysis (**C**); one-way ANOVA with post hoc Tukey’s (**D**); or two-sided unpaired Student’s t-test (**E, F**). P-values are shown in the figure*.

When compared with visceral nociceptor neurons, *Il6*, *Trpa1, Trpv1,* and *Npy1r* were also found to be significantly enriched in airway nociceptor neurons (**Fig. 1C-D**). Calcium imaging of JNC neurons showed that while capsaicin sensitivity was similar across neuronal subpopulations, JT010-mediated TRPA1 activation was drastically increased (2.5-fold) in airway neurons (DiD^+^) compared to other visceral neurons (**Fig. 1E, SF. 1D-E**). Using JNC neurons from NPY1R^Cre^::tdTomato^fl/wt^ mice, we also confirmed NPY1R enrichment in the airway population (**Fig. 1F**).

We then mapped out the airway nociceptor neuron subset within the JNC neuron populations using an in-silico analysis of Prescott and colleagues scRNAseq data^1^. We reanalyzed these data using *Vglut2, Scn10a* and *Scn1a* to define nociceptor neuron clusters and low-threshold sensory neurons^2^, and *Phox2b* and *Prdm12* to differentiate between jugular and nodose neurons (**Fig. 1G**, **SF. 2A-D**) We found that the marker genes of one cluster, coined *Nodose Nociceptor 8 (NN8)*, were highly enriched (Normalized Enrichment Score (NES)=2.2) in our airway nociceptor neurons sequencing (**Fig. 1G**).

NN8 appears to exclusively innervate the airways. Several of NN8 markers (*Trpv1, Trpa1*; **Fig. 1G**) are known drivers of neurogenic inflammation in the airways^40,47–50^, while this cluster also co-expresses *Npy1r* and *Il6* (**Fig. 1F, SF. 2E-J**). *Kcng1*, the top marker of NN8, is virtually absent in visceral nociceptors (**Fig. 1D**). Additionally, we identified three other populations of sensory neurons preferentially innervating the airways (**Fig. 1G)**, including a subset that we assume to be cough-inducing mechanoreceptors (NN7; **ST 1**). An in-silico analysis of Zhao et al.^7^ single cell projection-seq data validated this result (**SF. 3A-H**). Thus, these data confirm that one specific lung-innervating neuron subtype co-express *Scn10a* (gene encoding for Na_V_1.8), *Npy1r, Kcng1, Trpa1* and *Il13ra1* (**SF. 3C-H**).

### Airway nociceptors undergo transcriptional reprogramming in asthma

Next, we sought to test whether the airway nociceptor neuron transcriptome is impacted during allergic airway inflammation (AAI). Retrograde tracer-exposed nociceptor reporter mice (Na_V_1.8^cre^::tdTomato^fl/wt^) underwent the classic ovalbumin (OVA) model of asthma^18^. OVA-exposed mice showed significant airway inflammation characterized by eosinophilic infiltration in the bronchoalveolar fluid (**Fig. 2A**) but had a similar number of back-labeled airway nociceptor neurons (**SF. 4A**). Simultaneously, we evaluate how mice exposed to OVA modulate the influx of immune cells into the BALF of allergic mice by using single-cell RNA sequencing on flow cytometry-sorted lung CD45^+^ cells (**Fig. 2B**). The various cell types were identified using standard markers of lung immune cells, and include including B cells, T cells, Granulocytes (neutrophils and eosinophils), NK cells, Basophils, and Antigen Presenting Cells (APC, which include Macrophages and Dendritic cells) endothelial cells and alveolar macrophages (**SF. 4B-C**). DEGs detailed in the **ST. 2**, indicate significant airway inflammation with drastic changes in lung immune cells gene expression profiles.

**Figure 2.**
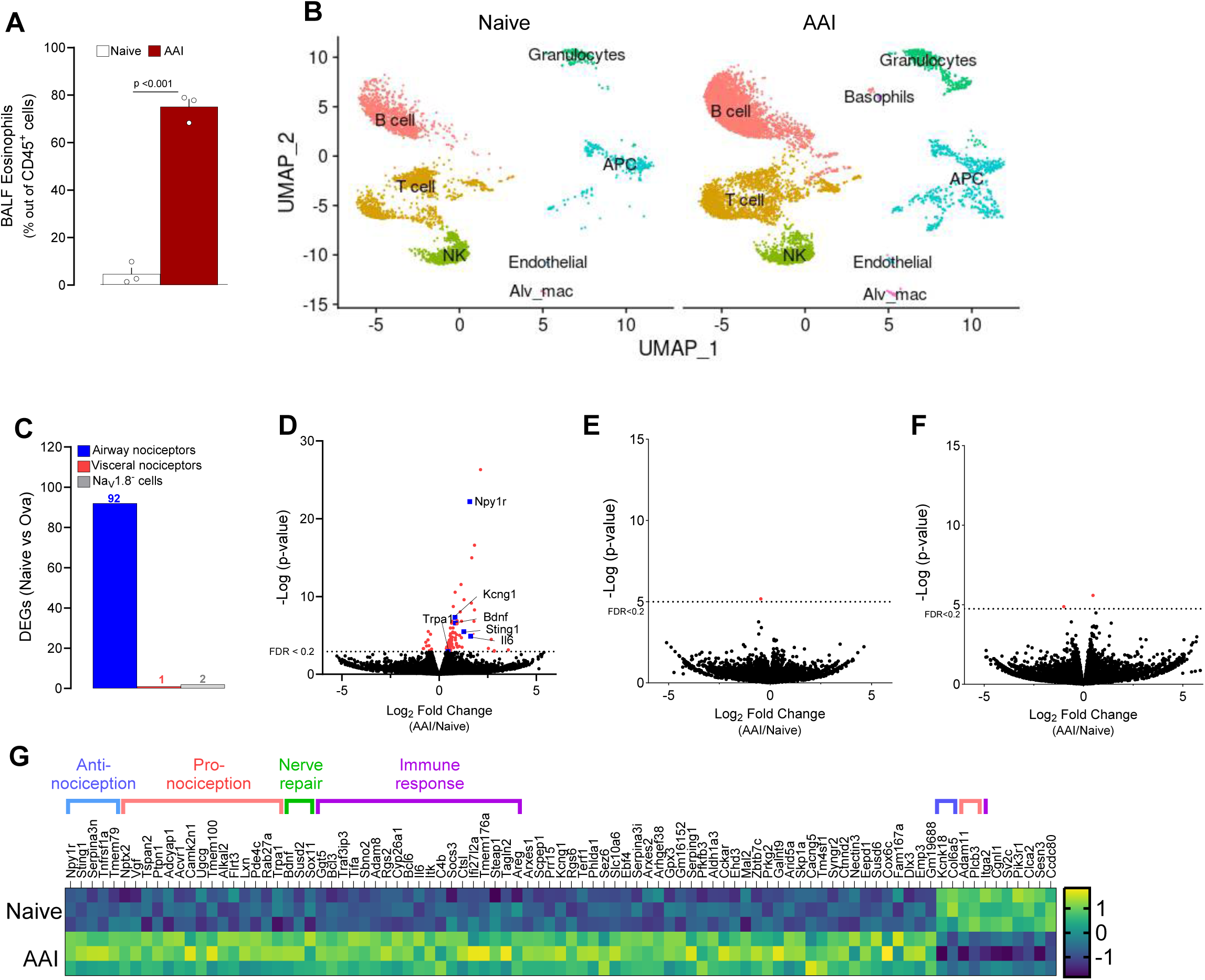
Allergic airway inflammation reprograms airway nociceptors transcriptome. (**A-B**) 8-week-old male and female nociceptor neuron reporter (Nav1.8^cre^::tdTomato^fl/wt^) mice underwent the ovalbumin mouse models of asthma. Allergic inflammation was induced in mice by an initial sensitization to ovalbumin (OVA) (i.p. day 0 and 7) followed by inhaled OVA challenges (days 14-17). Analysis of Bronchoalveolar lavage fluid showed significant airway inflammation characterized by a significant eosinophilic (CD45^+^CD11C^low^SiglecF^Hi^) infiltration (**A**). In other groups of mice, lungs were collected, and CD45^+^ cells were purified using FACS and subsequently analyzed through single-cell RNA sequencing. UMAPs demonstrated a change in the number and polarization of virtually all types of immune cells in the BALF of ovalbumin-exposed mice, including B cells, T cells, Granulocytes, NK cells, Basophils, and Antigen Presenting Cells (APC, which include Macrophages and Dendritic cells). **(B)**. **(C-G)** To identify airway-innervating nociceptor neurons, the mice were injected intranasally with the retrograde tracer DiD’ (200 µM) (days 2-4). One day after the last allergen challenge, the JNC ganglia were isolated and dissociated. Airway-innervating nociceptor neurons (Na_V_1.8^+^DiD^+^), visceral nociceptors (Na_V_1.8^+^DiD^-^) and Na_V_1.8^-^ cells were purified by flow cytometry and their RNA sequenced. Differentially expressed genes were virtually only observed in airway-innervating nociceptor neurons (**C**). Pairwise comparison shows differentially expressed transcripts in airway-innervating nociceptor neurons between naive and AAI conditions, shown as a volcano plot (adjusted p-value <0.2 in red; **D**), but limited changes in visceral nociceptors and Nav1.8-cells (**E** and **F** respectively). The significant DEGs observed in airway innervating neurons are displayed as a heatmap (**G**) Among others, *Il6, Sting1, Npy1r*, and *Bdnf* were overexpressed in airway-innervating nociceptor neurons (**D**, **G**). *Data are shown as mean ± S.E.M (**A;C**), as a Umap (**B**), as a volcano plot displaying DESeq2 normalized count fold change and nominal p-values for each gene **(D, E, F),** or a heatmap displaying the z-score of DESeq2 normalized counts **(G).** N are as follows: **a, c, d, e, f, g**: n=3 biological replicates (4 mice per sample), **b**, n=1 (2 mice per sample). P-values were determined by a two-sided unpaired Student’s t-test (**A**) DESeq2 analysis (nominal p-value, **D, E, F)**. P-values are shown in the figure*.

As previously defined (**Fig. 1A**), the JNC neurons were isolated, and airway nociceptor neurons (Na_V_1.8^+^DiD^+^), visceral nociceptor neurons (Na_V_1.8^+^DiD^-^) and Na_V_1.8^-^ cells were purified by flow cytometry and RNA-sequenced. DESeq2 analysis revealed that AAI significantly affected the expression of 92 genes in airway nociceptor neurons (**Fig. 2C, D**), compared with only one and two genes in the visceral nociceptor neurons and Na_V_1.8^-^ cell populations, respectively (**Fig. 2C, E-F**). Seventeen of these upregulated genes in airway neurons had previously been associated with increased pain or nociceptors activity, while six were observed to dampen pain sensation (**Fig. 2G**, **ST. 3**). Within the airway nociceptor neurons, AAI notably increased the expression of *Bdnf, Il6, Kcng1*, *Npy1r*, *Sting1,* and *Trpa1* (**Fig. 2D, G**).

### Cytokines trigger gene expression changes in nociceptors

Since the transcriptome changes induced by AAI are restricted to airway nociceptors and are virtually absent in other visceral nociceptors and Na_V_1.8^-^ cells, these variations are likely triggered by mediators in the airways detected by peripheral nerve endings. To identify these neuromodulators, we exposed (24h) cultured JNC neurons to various allergy driving cytokines, inflammatory lipids, and neurotrophins, and used transcription changes to *Npy1r*, *Sting1*, *Bdnf*, and *Il6* as proxies for an AAI-like signature (**Fig. 2D**). This screening approach revealed that IL-4 and IL-13 triggered the overexpression of *Npy1r* (**Fig. 3A)**, TNF-α and IL-1β induced *Il6* (**Fig. 3B**)*, Bdnf* was triggered by the neurotrophin itself **(Fig. 3C)**, while *Sting1* was induced by IL-1β, BDNF, TNF-α and IgE mixed with its cognate antigen ovalbumin **(Fig. 3D)**. The other cytokines and neurotrophins tested did not impact the expression of the tested genes.

**Figure 3.**
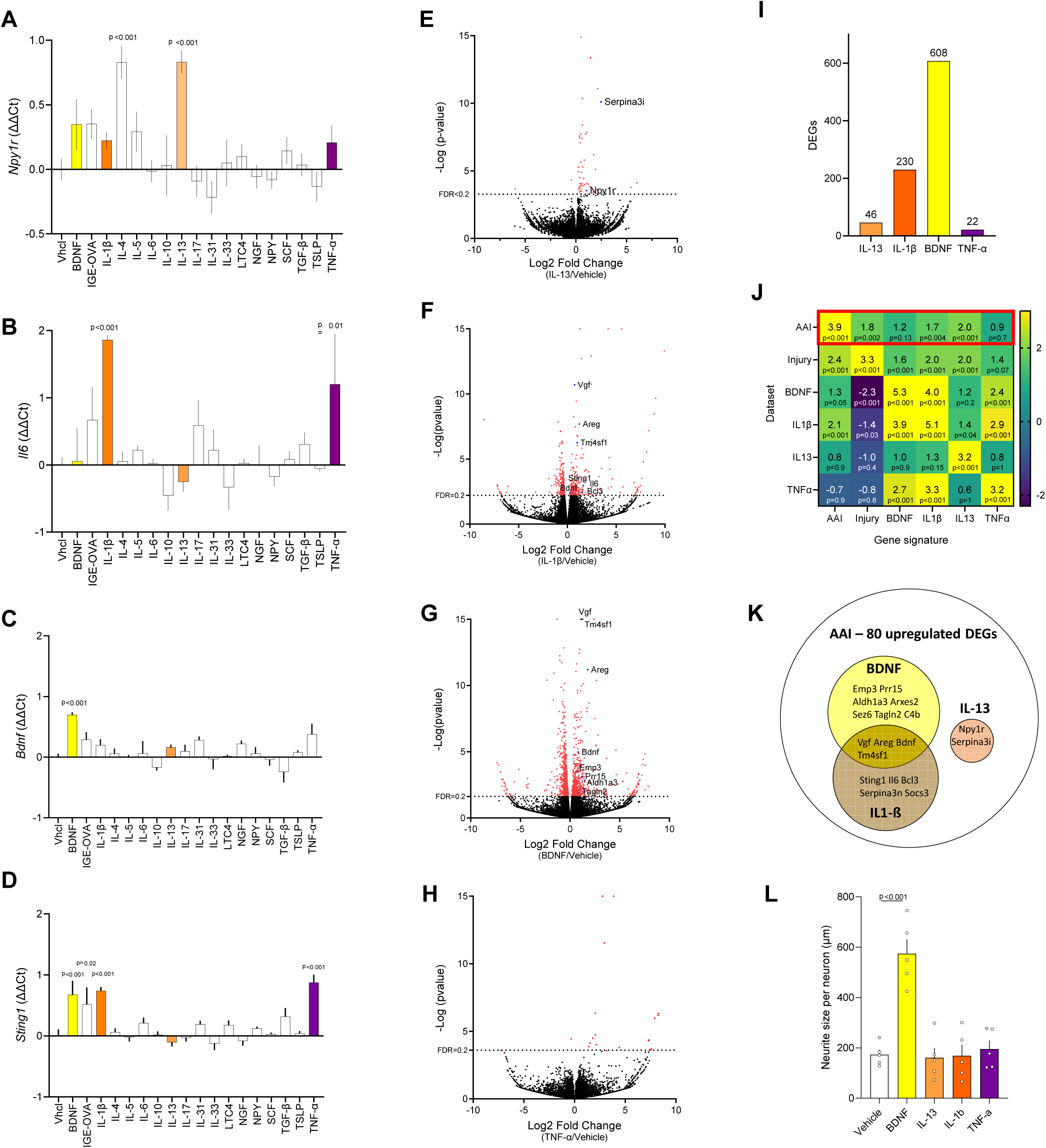
Screening of the cytokines driving AAI-induced nociceptor reprogramming. (**A-D**) 8-week-old male and female naïve C57BL6 mice JNC neurons were cultured and exposed (24h) to various inflammatory mediators. Changes in *Npy1r* (**A**), *Il6* (**B**), *Bdnf* (**C**), and *Sting1* (**D**) transcript expression were assessed by qPCR. In comparison to the vehicle, IL-4 and IL-13 (100 ng/mL) increased JNC neurons’ expression of *Npy1r.* IL-1β and TNF-α (100 ng/mL) increased the expression of *Il6*. BDNF induced its own overexpression while IL-1β (100 ng/mL), IgE-OVA (10 ug/mL), BDNF (50 ng/mL) and TNF-α (100 ng/mL) increased the expression of *Sting1*. (**E-I**) 8-week-old male and female naïve nociceptor neuron reporter (Na_V_1.8^cre^::tdTomato^fl/wt^) mice JNC neurons were cultured (24h) in the presence of vehicle, IL-13 (**E**), IL-1β (**F**), BDNF (**G**), or TNF-α (**H**). The nociceptor neurons (tdTomato^+^) were FACS-purified, and their RNA sequenced. Pairwise comparison showed differentially expressed genes induced by these cytokines (adjusted p-value <0.2 in red) and the genes in common with the AAI signature are highlighted in blue. IL-13, IL-1β, BDNF, and TNF-α respectively induced the overexpression of 46, 230, 608 and 22 genes (**I**). **(J)** GSEA analyses were performed to compare gene signatures (i.e., overexpressed DEGs) induced by AAI, nerve injury^53^, BDNF, IL-1β, IL-13 and TNF-α to their respective whole datasets. The gene signatures induced by nerve injury, IL-13, IL-1β and BDNF were enriched in JNC neurons from mice with AAI (**J**). (**K**) The common DEG induced by cytokines and AAI are depicted in a Venn diagram. IL-1β and BDNF signatures had similarities, while IL-13 induced two specific AAI genes *Npy1r* and *Serpina3i* (**K**). (**L**) Nociceptor neuron reporter (Na_V_1.8^cre^::tdTomato^fl/wt^) mice JNC neurons were cultured (24h) in the presence of vehicle, IL-13, IL-1β, BDNF, or TNF-α. BDNF, but not cytokines, promoted neurite growth (**L**). *Data are shown as mean ± S.E.M (**A-D, L**), as a volcano plot displaying DESeq2 normalized count fold change and nominal p-values for each gene (**E-H**) as number of DEGs (**I**), as heatmap displaying normalized enrichment score (NES) (**J**) or as a Venn diagram (**K**). N are as follows: **A-D**: n=3–4 cultures from different mice per group, **E-H:** n=3-4 cultures from different mice per group, **L**: n=6 culture dishes per group. P-values were determined by one-way ANOVA with post hoc Dunnett’s (**A-D, L**) or DESeq2 analysis (**E-H**), or GSEA analysis (**J**). P-values are shown in the figure or indicated by * for p ≤ 0.05; ** for p ≤ 0.01; *** for p ≤ 0.001*.

Cytokines and neurotrophins are known to initiate transcription programs in their target cells^51,52^. We then exposed cultures of JNC from Na_V_1.8^cre^::tdTomato^fl/wt^ reporter mice to IL-13, IL-1β, BDNF and TNF-α (24h), before purification of the nociceptor neurons by flow cytometry (tdTomato^+^) and RNA-sequencing. DEGs were triggered in the four different conditions **(Fig. 3E-I, ST. 4)**, with the most drastic changes induced by IL-1β and BDNF.

To compare reprogramming induced by AAI and the tested cytokines, a gene set enrichment analysis (GSEA) was performed using the most significantly overexpressed genes (FDR<0.2) in all tested conditions. These data revealed that the gene sets elevated in cultured JNC neurons exposed to IL-1β, IL-13, and BDNF, as well as the ones previously identified in injured neurons^53^, were enriched in JNC airway nociceptors of AAI mice (**Fig. 3J, ST. 4**). IL-13 signature showed the strongest enrichment (NES=2.0). Specifically, 11 DEGs identified in AAI neurons were then also induced *in vitro* by BDNF, and 9 genes induced by IL-1β, among which 4 were induced by these two proteins (**Fig. 3K)**. IL-13 induced 2 genes identified in AAI, *Npy1r* and *Serpina3i* (**Fig. 3K)**. Morphologically, BDNF influenced neurite outgrowth in cultured JNC nociceptors, whereas the tested cytokines did not have any effect (**Fig. 3L**). These results highlight the similarities and specificities between transcription programs inducible in nociceptor neurons, and suggest that IL-1β, BDNF and IL-13 signaling pathways are complementary to induce the AAI signature in our mouse model of asthma.

### IL-13 reprograms nociceptors through JAK/STAT6

While IL-1β and BDNF effects on nociceptors has been thoroughly investigated^6,50,54–57^, the literature regarding IL-13 is scarcer^14,58^, and this interaction was not yet reported in vagal sensory neurons. We then aimed to delineate the mechanisms by which IL-13, a type 2 specific cytokine, activates nociceptor neurons and changes their transcriptome. IL-13 (24h) significantly affected the expression of 47 genes, including *Npy1r* (**Fig. 3B, C, Fig. 4A, ST. 5**). Since both IL-4 and IL-13 and had comparable effects *in vitro* (**Fig. 3A),** we used our transcriptomic data to identify which IL-13 and IL-4 receptors are expressed by nociceptor neurons and found that only the IL4RII subunits *Il4ra* and *Il13ra1* are detected in these cells (**SF. 5A, ST. 6**). *Il13ra1* expression was higher in nociceptors than in Nav1.8^-^ cells (**SF. 5A**) and was also found to be expressed in the inflammatory airway nociceptor cluster NN8 (**Fig. 1E**; **SF. 2I**), along with the signaling mediator *Stat6* (**SF. 2J)**.

**Figure 4.**
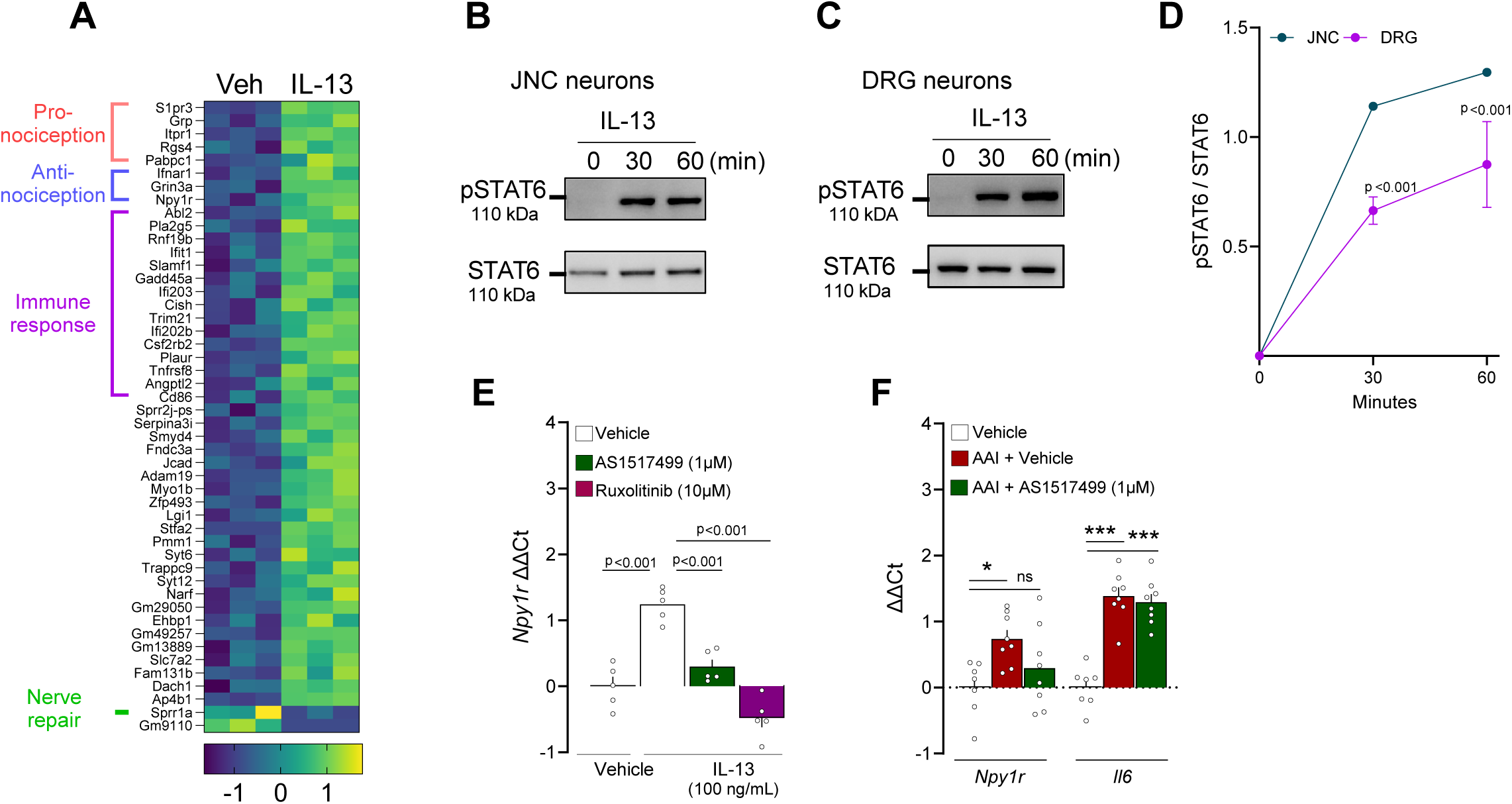
IL-13 reprograms nociceptor neurons. (**A**) Pairwise comparison shows 48 differentially expressed genes between the vehicle and IL-13 treated neurons, shown as a heatmap with putative gene function (see Supplementary table 4) (**A**). (**B-D**) Naive 8-week-old male and female C57BL6 mice JNC or DRG neurons were cultured and exposed (30– 60 minutes) to IL-13 (100 ng/mL) or vehicle. Changes to pSTAT6 were analyzed by Western blot. IL-13 time-dependently increased the pSTAT6/STAT6 ratio in JNC (**B, D**) and DRG (**C, D**) neurons. **(E)** In the presence or absence of the STAT6 inhibitor AS1517499 (1 µM) or the JAK1/2 inhibitor ruxolitinib (10 µM), naive 8-week-old male and female C57BL6 mice JNC neurons were cultured (24h) with IL-13 (100 ng/mL) and changes to *Npy1r* transcript expression was analyzed by qPCR. *Npy1r* overexpression induced by IL-13 was blocked by both AS1517499 and ruxolitinib (**E**). (**F**) 8-week-old female nociceptor neuron reporter (Na_V_1.8^cre^::tdTomato^fl/wt^) mice underwent the ovalbumin mouse models of asthma. Allergic airway inflammation was induced in mice by an initial sensitization to ovalbumin (OVA) (i.p. day 0 and 7) followed by intranasal instillation with OVA mixed with STAT6 inhibitor AS1517499 (150ug/50ul) or vehicle (days 14-17). One day after the last allergen challenge, mice were sacrificed, and JNC nociceptor neurons (tdTomato^+^) purified by flow cytometry for RNA extraction and RT-qPCR. OVA-exposed mice nociceptors showed increased expression of *Npy1r* and *Il6*. *Npy1r* overexpression was prevented by AS1517499 (**F**). *Data are shown as a heatmap displaying the z-score of DESeq2 normalized counts **(A),** as Western blots **(B-C),** as mean ± S.E.M (**D, F**), or as box (25th–75th percentile) and whisker (min-to-max) plot (**E**). N are as follows: **A:** n=3 cultures from different mice per group, **B-D**: n=1 JNC culture and n=3 DRG cultures from different mice, **E**: n=5 cultures from different mice per group, **F**: n=7-8 mice per group. P-values were determined by one-way ANOVA with post hoc Dunnett’s (**D-F**). P-values are shown in the figure or indicated by * for p ≤ 0.05; ** for p ≤ 0.01; *** for p ≤ 0.001*.

Since IL4RII is a shared receptor for IL-13 and IL-4 that signals via JAK1/2 and STAT6 to regulate immune cells’ transcription^59^, we tested whether a similar mechanism was at play in nociceptor neurons. We found that IL-13 (30min) triggered STAT6 phosphorylation in JNC and DRG neuron cultures (**Fig. 4B-D**). In addition, IL-13-mediated induction of *Npy1r* was prevented by the STAT6 inhibitor AS1517499 as well as by the JAK1/2 inhibitor ruxolitinib (**Fig. 4E**). In the tested conditions, IL-13 did not induce calcium flux in nociceptor neurons (**SF. 5B**). The effect of IL-13 on *Npy1r* expression in JNC nociceptors was also observed in cultured DRG neurons (**SF. 5C**). To see if this pathway is responsible for *Npy1r* overexpression *in vivo,* we treated AAI mice with intranasal instillation of AS1517499 (150µg/50µL). AS1517499 prevented *Npy1r* induction in nociceptor neurons without affecting *Il6* overexpression, confirming that the AAI molecular profile depends in part on pSTAT6 activity **(Fig. 4F)**.

### Sympathetic neurons and APCs produce NPY in the airways during asthma

*Npy1r* thus appears as a major marker of AAI inflammation in nociceptor neurons. Since the optogenetic activation of NPY1R^+^ JNC neurons trigger expiratory reflexes in mice^1^ and NPY1R activation impacts pain and itching reflexes^60–68^, we sought to test whether Neuropeptide Y (NPY)/NPY1R interaction could occur in the lung. Along with airway inflammation (**Fig. 5A-B**), allergen challenges (day 18) significantly increased the release of NPY in BALF (**Fig. 5C**) and serum (**Fig. 5D**). Of note, NPY levels were normal upon allergen sensitization (day 1 and 7), and while immune cells were still elevated, the neuropeptide concentration returned to baseline during the resolution phase (day 21; **Fig. 5C-D).** A similar pattern was observed for BALF IL-13 level (**Fig. 5C-D**). Concomitantly, *Npy1r* overexpression in JNC nociceptors also transiently peaked upon allergen challenges (**Fig. 5E**). We then tested whether these changes were specific to OVA-induced airway inflammation model and found a similar rise in BALF NPY in the house dust mite (HDM) model of asthma (**Fig. 5F**).

**Figure 5.**
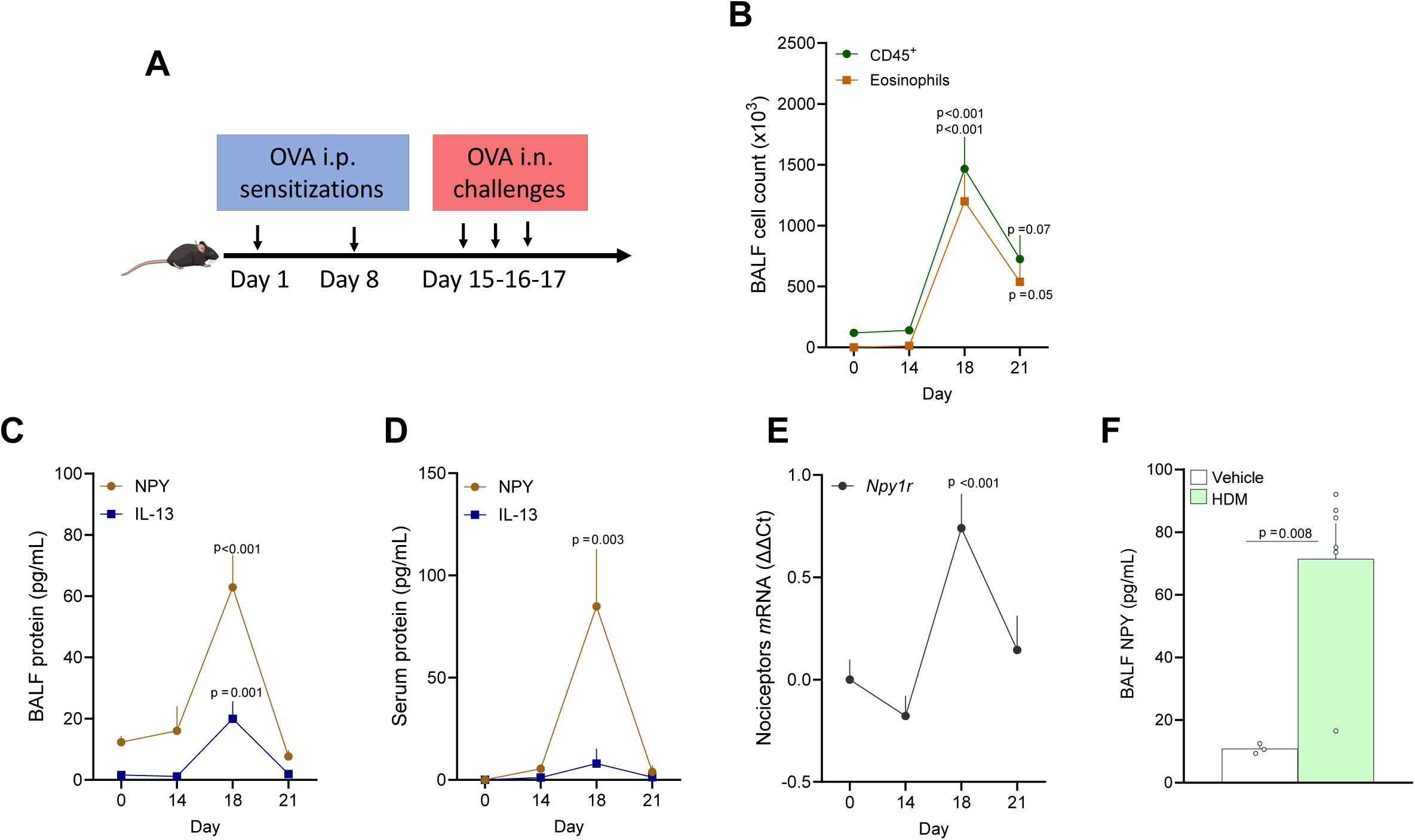
NPY is released in airways during allergic airway inflammation. **(A)** Schematic of the AAI protocol. 8-week-old female C57BL6 mice underwent the ovalbumin mouse models of asthma. Allergic airway inflammation was induced in mice by an initial sensitization to ovalbumin (OVA) (i.p. day 0 and 7) followed by inhaled OVA challenges (i.n. days 15-17). **(B-D)** 8-week-old female C57BL6 mice underwent the ovalbumin mouse models of asthma. BALF, serum and lung were harvested at different time points (days 0, 14, 18, and 21) and the levels of inflammatory mediators were analyzed by ELISAs and qPCR. OVA-exposed mice showed significant airway inflammation characterized by leukocytes (CD45^+^) and eosinophil (CD45^+^CD11C^low^SiglecF^Hi^) infiltration on day 18 (**F**). Along with this rise in airway inflammation, we found an increase in BALF (**G**) and serum (**H**) NPY, while lung *Npy* expression was also increased (**I**). **(E)** JNC ganglia were harvested from OVA-exposed nociceptor reporter mice (TRPV1^cre^::tdTomato^fl/wt^) at different time points, TRPV1^+^ neurons were purified (tdTomato^+^) by flow cytometry, and changes in transcript expression were measured by qPCR. At the peak of inflammation (day 18), we found a transient increase in *Npy1r* expression (**J**). **(F)** To induce another model of allergic airway inflammation, 8-weeks-old female C57BL6 mice were challenged (day 1-5 and 8–10) with house dust mite (HDM; 20μg/50μL, i.n.). The mice were sacrificed on day 11, their BALF harvested, and cell free supernatant analyzed by ELISA. HDM-exposed mice showed a significant increase in BALF NPY level (**F**). *Data are shown as experiment schematics (**A**), or as mean ± S.E.M (**B-F**). N are as follows: **B-D**: n=6-7 mice per group, **E**: n=4-10 mice per group, **F:** n= 3-6 mice per group. P-values were determined by one-way ANOVA with post hoc Dunnett’s (**B-E,** comparison to day 0) or two-sided unpaired Student’s t-test (**F**). P-values are shown in the figure or indicated by * for p ≤ 0.05; ** for p ≤ 0.01; *** for p ≤ 0.001*.

Using lung cryosections and immunostainings, we observed a strong and specific expression of NPY in nerve fibers often located around the bronchi (**Fig. 6A**). Staining of lung innervating peripheral neuron ganglia showed that 40% of sympathetic neurons in the stellate ganglia (SG) express NPY (**Fig. 6B-D**) which is in sharp contrast with virtually no expression in the JNC neurons (**Fig. 6B-D**). Using triple-labeling, we found that NPY-expressing neurons around the bronchi colocalized with the sympathetic neuron marker tyrosine hydroxylase (TH; **Fig. 6E-F**). While in proximity, the NPY-expressing neuron fibers were mostly distinct from Na_V_1.8 positive nociceptor nerve endings (**Fig. 6A, E, F**). We confirmed that NPY positive nerve fibers in the lung were of sympathetic and not sensory origin by performing a chemical sympathectomy with 6-OHDA (**Fig. 6G-H**). 6-OHDA treatment completely abolished NPY and TH expression in the lung but did not affect the presence of CGRP^+^ nociceptor neurons (**Fig. 6G-H**).

**Figure 6.**
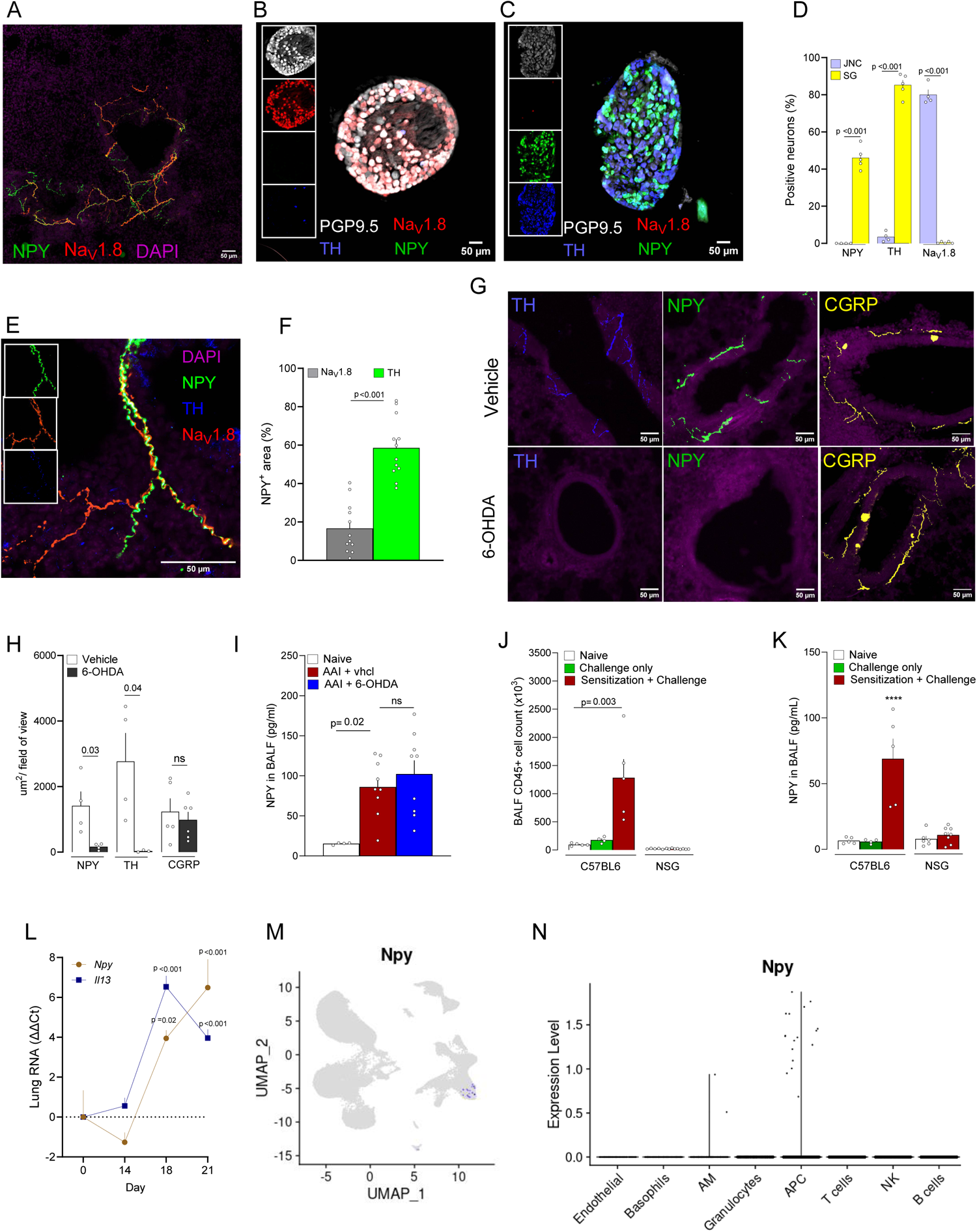
NPY is expressed by sympathetic neurons and inflammatory macrophages in the lung. **(A-F)** Lung (**A**), jugular nodose complex ganglia (JNC; **B**) and stellate ganglia (SG; **C**) were harvested from naïve 8-weeks-old male and female nociceptor neuron reporter (Na_V_1.8^cre^::tdTomato^fl/wt^) mice. The tissues were cryosectioned, and the source of NPY assessed by immunofluorescence. In the lung, NPY (green) and Na_V_1.8 (tdTomato, red) were expressed in nerve fibers around the bronchi **(A).** NPY was not expressed in the JNC (**B, D**). In the stellate ganglia (SG), NPY was strongly expressed in TH^+^ sympathetic neurons (**C, D**). PGP9.5 (white) was used to define JNC and SG neurons (**B, C, D**). While sympathetic and sensory fibers were often found in proximity, NPY (green) mostly colocalized with the sympathetic neuron marker tyrosine hydroxylase (TH; blue) rather than with Na_V_1.8 (tdTomato; red) nociceptor fibers **(E-F).** NPY and TH were absent in lung cryosections from mice with chemical sympathectomy induced by OHDA **(G-H)**, while sensory nerve fibers (CGRP+) were not affected **(G-H)**. **(I)** NPY is still elevated in bronchoalveolar lavage fluid of AAI mice with chemical sympathectomy **(I)**. **(J-K)** Immunodeficient mice (NSG) showed absence of immune cell infiltration in the airways following OVA protocol **(J)**. While NPY was released in the BALF of wild type mice with AAI, it was absent in the airways of immunodeficient mice (NSG) **(K)**. Additionally, OVA challenge without previous sensitization neither induced immune response (**J**) nor NPY release in the airways (**K**). **(L-N)** NPY was elevated in the RNA of mice at the peak of inflammation of the AAI protocol, (day18), and persisted 3 days later (day 21) **(L)**. Single-cell RNA sequencing of FACS-purified CD45^+^ cells revealed selective expression of *Npy* in a subset of antigen-presenting cells (APC) and macrophages within the bronchoalveolar lavage fluid (BALF) of ovalbumin-exposed mice **(M-N)**. *Data are shown as immunostained tissue, scale bar 50 μm (**A-C, E, G**), or as mean ± S.E.M (**D, F, H-L**), or Seurat normalized gene expression (**M, N**). N are as follows: **D**: n=4–5 mice per group, **F:** n=12 field of views from 4 different mice, **D**: n=4–5 mice per group, **I**: n=4–10 mice per group, **J-K**: n=4–10 mice per group, **L**: n=6–7 mice per group. P-values were determined by a two-sided unpaired Student’s t-test (**D, F, H**) or by one-way ANOVA with post hoc Dunnett’s (**I, J, K**). P-values are shown in the figure*.

While it might seem logical to conclude that sensory and sympathetic fibers interact via NPY/NPY1R due to their close proximity, 6-OHDA-induced sympathectomy surprisingly did not affect NPY release in the airways of asthmatic mice (**Fig. 6I**). However, in the AAI model, NOD scid gamma (NSG) immunodeficient mice exhibited no increase in BALF NPY levels in the airways (**Fig. 6J-K**), suggesting that immune cells might also be a source of NPY. Moreover, both OVA sensitization and challenge were necessary to trigger NPY release in the airways (**Fig. 6K**), confirming that the full adaptive immune response to the allergen is essential. Supporting this hypothesis, we observed an increase in *Npy1r* RNA expression in the entire lung from mice with AAI (**Fig. 6L**), indicating NPY production by non-neuronal cells. Through our single-cell RNA-seq of AAI lung immune cells (**Fig. 2B**), we identified *Npy* expression exclusively in a subtype of antigen-presenting cells (APC) (**Fig. 7M-N**), which bear M2 macrophages markers such as *Fcgr1, Ccr5, Cd63, Mrc1, Msr1, Ccl24* **(SF. 6A**). This finding aligns with a prior study that reported NPY expression in phagocytes in a mouse influenza model^69^. We conclude that, although sympathetic neurons might interact with sensory neurons due to their proximity, the primary drivers of this interaction during AAI are likely NPY-producing M2 macrophages.

**Figure 7.**
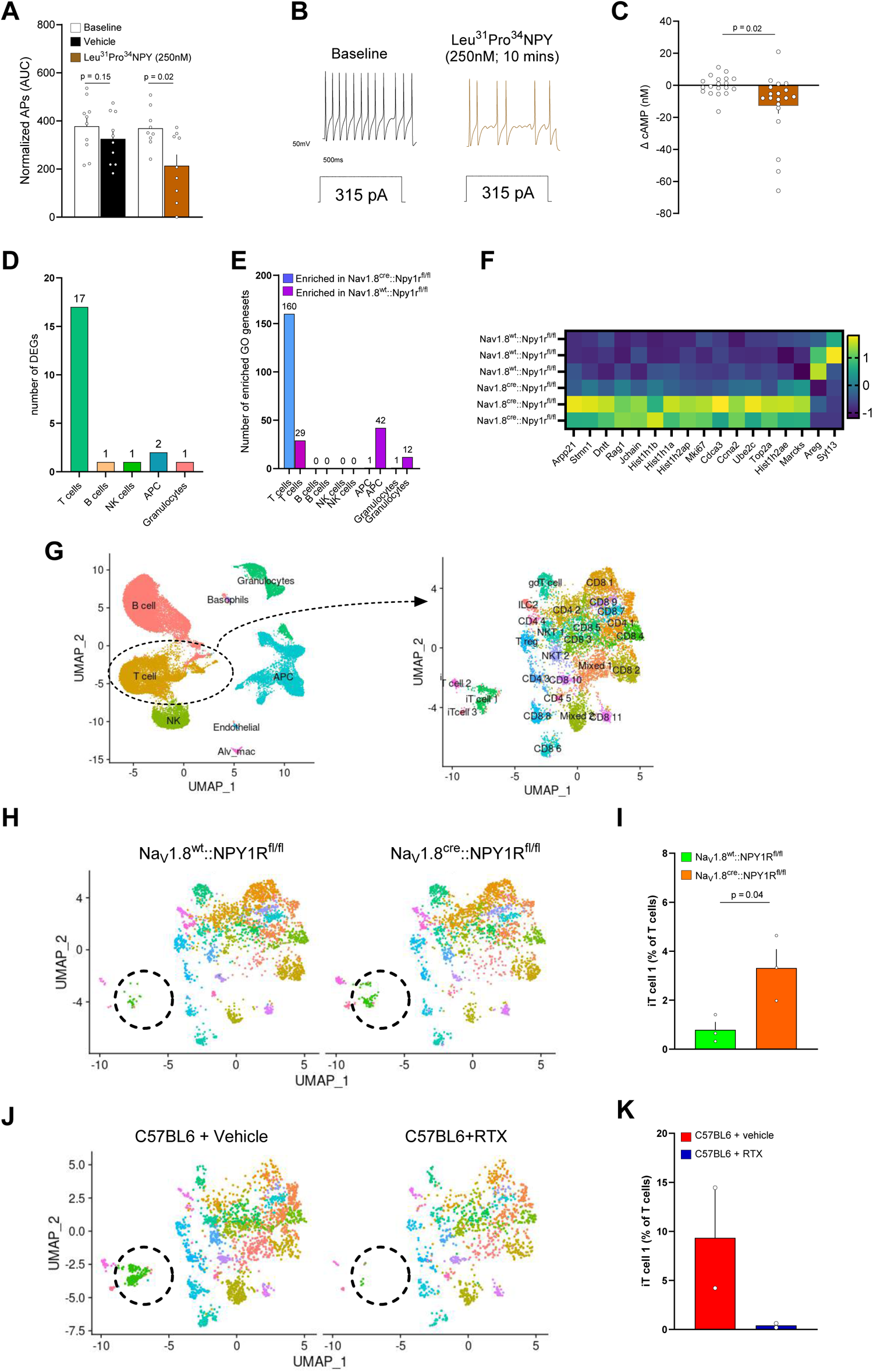
NPY1R blunts JNC nociceptors excitability. **(A-C)** 8-week-old male and female NPY1R reporter (NPY1R^cre^::tdTomato^fl/wt^) mice were sacrificed and their JNC neurons harvested and cultured (16 hours). Whole cell patch clamp electrophysiology was performed on the NPY1R^+^ nociceptor neurons. A current clamp was applied while the neurons’ membrane potential was recorded before and after exposing (10 min) the cell to Leu^31^Pro^34^NPY (250 nM) or its vehicle. The number of action potentials was counted for each current stimulation, and the areas under the curve was calculated and plotted **(A)**. A representative trace of neuronal response to 315pA stimulation before and after Leu^31^Pro^34^NPY is displayed in **(B)**. While the vehicle had little to no effect on neuronal excitability, Leu^31^Pro^34^NPY reduced the number of action potentials in response to current stimulation in NPY1R^+^ (tdTomato) neurons **(A, B).** **(C)** 8-week-old C57BL6 male and female mice were sacrificed and their JNC neurons harvested and cultured (16 hours). The neurons were exposed to Leu^31^Pro^34^NPY (250 nM) or vehicle for 30 minutes in presence of phosphodiesterase inhibitors. The cells were then lysed, and the cAMP concentration assessed by enzymatic assay. Leu^31^Pro^34^NPY significantly reduced cAMP concentration **(C)**. (**D-F**) Eight-week-old female mice with a nociceptor neuron-specific NPY1R conditional knockout (Na_V_1.8^cre^::NPY1R^fl/fl^) and their littermate controls (Na_V_1.8^wt^::NPY1Rf^l/fl^) were subjected to the ovalbumin-induced asthma model. On day 18, lungs were harvested, and CD45^+^ cells were purified by flow cytometry before undergoing analysis by single-cell RNA sequencing. Gene expression was averaged for each cell type in each replicate to conduct pseudobulk analysis using DESeq2 (**D**) and gene set enrichment analysis (**E**). Na_V_1.8^cre^::NPY1R^fl/fl^ mice exhibited exacerbated allergic airway inflammation, with T-cells displaying the most significant differences in terms of the number of differentially expressed genes (**D**; False Discovery Rate < 0.2) and enriched Gene Ontology gene sets (**E**; False Discovery Rate < 0.2). The average expression of all T-cell DEGs is displayed in a heatmap (**F**). *Complete DESeq2 analysis (**D**) and GO gene sets (**E**) are detailed in Supplementary Tables 8 and 9*. (**G-K**) The T cells isolated from the single-cell RNA-seq dataset of AAI-affected lungs were subclustered (**G**). In the UMAP projections, the cluster named immature T cells 1 (iTcells 1) appears distinct between the NPY1R conditional knockout mice (Na_V_1.8^cre^::NPY1R^fl/fl^) and the littermate control mice (Na_V_1.8^wt^::NPY1R^fl/fl^) (**G**). Quantification reveals a significantly higher number of these cells in the Na_V_1.8^cre^::NPY1R^fl/fl^ mice (**H**). In contrast, iTcells 1 were less abundant in nociceptor neurons ablated (RTX-treated) mice compared to vehicle control (**I-J**). *Data are shown as traces of membrane potential for individual neurons (**B**), mean ± S.E.M **(A, C)**, or UMAP* ***(D)****. N are as follows: **A**: n=10 vehicle treated neurons and 9 Leu^31^Pro^34^NPY treated neurons, **C**: n=19 culture wells, **D-I:** n=3 pools of 2 mice per group, **J-K**: n=2 pools of 2 mice per group. P-values were determined by a two-sided unpaired Student’s t-test **(A, C, I, K),** Deseq2 analysis (**D**), GSEA analysis (**E**). P-values are shown in the figure*.

### NPY1R blunts vagal nociceptor excitability

NPY1R is a Gi-coupled receptor^70^ and the action of NPY on DRG nociceptor neurons promotes either analgesia or noxious hypersensitivity^60–68^. Thus, A significant body of literature shows that NPY-NPY1R signaling contributes to the pathology of bronchial asthma and airway hyperresponsiveness in mouse models of allergic airway inflammation^71,72^. Additionally, NPY polymorphisms have been linked to an increased risk of asthma in overweight individuals^73^. Given that both *Npy1r* and NPY levels were elevated during allergic airway inflammation, we set out to address how NPY-NPY1R modulates cultured JNC nociceptor neuron sensitivity. To do so, we used whole cell patch clamp recording of NPY1R^+^ JNC nociceptor neurons (NPY1R^cre^::tdTomato^fl/wt^). We measured electrical changes in response to the NPY1R-specific agonist Leu^31^Pro^34^NPY. We observed a significant reduction in the nociceptor neurons’ excitability (**Fig. 7A-B, SF. 7A-C**), as measured by a reduced number of action potentials in response to current injection. We also measured a reduced level of intracellular cAMP in cultured JNC neurons exposed to Leu^31^Pro^34^NPY (**Fig. 7C**). Both effects indicate a direct engagement of NPY1R on nociceptor neurons and a strong reduction in the electrical activity and response capacity of these neurons.

To comprehensively understand the impact of NPY1R expression on sensory neurons on the asthma phenotype, we generated mice with a conditional knockout of this receptor in nociceptors (Na_V_1.8^cre^::Npy1r^fl/fl^). First, we used flow cytometry to immunophenotype the BALF of OVA-exposed Na_V_1.8^cre^::Npy1r^fl/fl^ and its littermate control, and found similar immune cell numbers across both groups (**SF. 8A-B**). Next, we used single-cell RNA sequencing (scRNA-seq) along with a pseudo-bulk and DESeq2 analysis approach to map the entire airway immune cell landscape in mice that underwent the asthma induction protocol. In the absence of NPY1R, we observed changes in T-cell populations, including 17 differentially expressed genes (DEGs) and the modulation of 160 Gene Ontology (GO) gene sets (**Fig. 7D-F**, **ST. 7-9).** Looking at T-cell subpopulations in detail, changes were primarily observed in the *Rag1*+ immature T-cell 1 (iT-cell-1) cluster (**Fig. 7G-I, SF. 8C-G, ST. 10**). We noted an increased proportion of these cells in the conditional knockout (Na_V_1.8^cre^::Npy1r^fl/fl^) mice compared to their littermate control animals. Most notably, the chemical ablation (RTX-exposed mice) of nociceptor neurons had the opposite effect, completely eliminating the emergence of this T-cell cluster in the BALF of asthmatic mice (**Fig 7J-K)**.

## DISCUSSION

Nociceptor neurons shape host defense at mucosal barriers. They do so by detecting environmental danger, triggering an avoidance response, and by tuning immune responses. In the context of allergy, nociceptor neurons were found to amplify dermatitis^14,19,20,74^, conjunctivitis^28^ and airway inflammation^18,30,47,48,75,76^. In the lungs, these responses range from coughing and bronchoconstriction to mucus secretion and depending on the context, amplifying, or taming immunity. Such broad responses are made possible by the highly heterogenous nature of nociceptor neurons.

### Airway nociceptors

The exact neuronal subset involved in these responses remained to be defined. Zhao and colleagues^7^ posit that the variety of organs innervated by the vagus nerve explains in part the need for JNC sensory neuron heterogeneity. As such, airway sensory neurons^8,9,11,46^ were classified as i) low-threshold stretch-sensitive neurons (essential to the respiratory cycle); ii) mechanoreceptors (sensitive to punctate mechanical stimuli); and iii) high threshold thermosensitive and chemosensitive nociceptors (recruited in response to tissue injury, inflammation, noxious chemicals or temperatures).

Using a combination of lineage reporters, retrograde tracing and transcriptomic analysis, we revealed that JNC airway nociceptor neurons have a unique gene signature segregated from that of other visceral nociceptors. We identified a new class of *Kcng1*-expressing inflammatory nociceptors (*Trpa1^+^*, *Trpv1^+^*, *Il6^+^*, *Npy1r^+^, Il13ra1^+^*) that exclusively innervate the airways (NN8). This confirms the assumption of Kupari and colleagues^2^ that this neuron subtype (which they labeled as NG14) consists of pulmonary afferent unmyelinated neurons. Additionally, we found that the airways are preferentially innervated by a neuronal subset (*Kcnv1^+^*, *Piezo1^+^*, *Piezo2^+^*) reminiscent of cough mechanoreceptors (NN7, NG3 in Kupari et al.’s study) as well as by a subset of polymodal nociceptors (NN2), and an Na_V_1.8^low^ population possibly belonging to the low-threshold stretch-sensitive neurons^10,11^. These neuronal subtypes and their markers are listed in **supplementary table 1.**

### AAI reprogramming

We identified a drastic reprogramming of airway nociceptor neurons in response to allergic airway inflammation. Interestingly, the gene signature that we identified largely overlap with the one typically observed in injured nociceptor neurons. We can hypothesize that inflammation induces neuronal damage in the airways, which would explain such similarities. These changes are also reminiscent of those observed in LPS model of airway inflammation^77^.

While immune and glial cell activation or infiltration is sometimes reported in the spinal cord or DRG following peripheral inflammation or nerve injury^66,78–82^, this was not the case in the JNC of AAI mice. This implies that the neuro-immune interactions occur at the peripheral nerve ending level rather than in the ganglia. Additionally, the nociceptor neurons innervating other organs were shielded from the transcriptional changes we observed in airway neurons, which also involves that those changes are triggered by signals originating from the nerve ending in the airways.

### Cytokines reprograms nociceptor neurons

As we found that AAI reprogrammed airway nociceptor neurons’ transcriptome, we tested how various cytokines impact JNC neuron expression profiles. Interestingly, nociceptor neurons showed different gene signatures when exposed to IL-13/IL-4, IL-1β, TNF-α, or BDNF. Thus, it appears that the combination of signaling pathways induced by nerve ending damage, inflammatory cytokines and neurotrophins add up *in vivo* and result in the nociceptors AAI signature. These findings are also indicative of nociceptor neurons’ plasticity to various inflammatory conditions and subsequent context-dependent neuro-immune responses^34^.

IL-13 mimics some of the transcriptional changes observed during AAI, an effect that involves STAT6 phosphorylation and subsequent regulation of gene expression. In physiology, our findings suggest that IL-4/IL-13 released by airway T_H_2 and ILC2 cells are locally sensed by IL4RII-expressing vagal nerve endings. In turn, the intracellular signals are likely to be retrogradely transported to the soma to generate transcriptional changes. Such retrograde transport has been reported in nociceptors for STAT3^83–86^ and CREB^87,88^, and thus may also occur for STAT6 (**SF. 7**).

IL-4 and IL-13 were previously found to induce calcium flux in dorsal root ganglia nociceptor neurons and to trigger itching^14,58^. While we did not observe direct calcium flux in JNC nociceptor neurons exposed to IL-13, our data converge regarding the functional expression of IL4RII and JAK1/2 activation in nociceptor neurons^58,89^. IL-13/IL-4/JAK/STAT6 is a key signaling pathway essential to type 2 inflammation and allergies^90,91^, and strategies to target it have proven effective to treat atopic dermatitis, asthma and to prevent itching^92^. We can reason that the sensory relief observed in these patients may, in part, be due to the silencing of this pathway in nociceptor neurons.

### NPY, NPY1R, pain and allergy

Pain warns the organism of environmental dangers. Endogenous neuromodulatory mediators can either increase or decrease the organism’s perception. In the context of pain, the impact of NPY and NPY1R remains controversial. Nevertheless, a consensus has emerged as for NPY1R expression and antinociceptive effect in the central nervous system^60–63,65^.

The effect on primary afferent nociceptor neurons is less established and NPY exerts a complex influence on pain sensitivity and neuropeptide release. This duality is likely explained by nociceptor neurons’ co-expression of NPY1R and NPY2R, with the former dampening pain and inflammation while the latter exacerbates it^64,66–68^. Building on these findings, we presented the first set of data suggesting the impact of NPY-NPY1R on vagal neuron sensitivity. We discovered that NPY-NPY1R decreased JNC nociceptor neuron activity by decreasing the levels of cAMP and reducing action potential firing in response to electrical stimulation. cAMP has long been recognized as an intracellular messenger promoting nociceptor sensitization^93–98^, an effect mediated in part by PKA-induced phosphorylation of Na_V_1.8 channels^99^. NPY also fulfills numerous physiological functions, ranging from the control of hunger/feeding to energy homeostasis^100,101^, vasoconstriction^102^ and immunomodulation^103^. As we found to be the case in our OVA- and HDM-challenged mice, other studies showed elevated NPY levels in various rodent models of lung inflammation^71,72,104–106^ as well as in the plasma of elderly asthma patients^107^. Intriguingly, most of the NPY released appeared to originate from macrophages, rather than from sympathetic neurons.

Our single cell RNA sequencing data now reveal that NPY1R-expressing nociceptor amplify, the regulation and function of T cells, with implications for asthma, allergies, and broader inflammatory processes. *Arpp21* and *Stmn1* influence RNA splicing and microtubule dynamics, respectively, affecting T cell activation and migration crucial for allergic responses. *Dntt* and *Rag1* are pivotal in the development of diverse T-cell receptor repertoires, directly impacting sensitivity to allergens and the immune response’s adaptability. *Jchain,* though more closely associated with B cells, contributes to mucosal immunity relevant in asthma by mediating IgA and IgM transport. Histone genes like *Hist1h1b, Hist1h1a, Hist1h2ap,* and *Hist1h2ae* regulate gene expression through chromatin remodeling, influencing cytokine production in allergic inflammation. Cell cycle regulators such as *Mki67, Cdca3, Ccna2, Ube2c,* and *Top2a* affect T cell proliferation, a key event in the immune response to allergens. *Marck*s’ role in cell signaling and migration is critical for T cell trafficking to inflammation sites, while *Areg* is involved in tissue repair and remodeling in asthma. Lastly, *Syt13’s* potential impact on cytokine release through vesicular trafficking highlights the complex network of genes influencing T cell behavior and the pathophysiology of asthma, allergies, and inflammation, offering insights into potential therapeutic targets for these conditions. This supports the hypothesis that NPY1R reduces airway nociceptor activity, thereby diminishing their impact on the airway immune response in asthma. Knocking out NPY1R disrupts this regulatory pathway, leading to an outcome opposite to that of neuronal inactivation by resiniferatoxin. NPY was also reported to increase methacholine-induced bronchoconstriction^71,104^ while NPY1R^+^ vagal neurons were shown to trigger expiratory reflexes^1^. It would thus be of interest to assess whether its expression on vagal neurons is involved in cough and bronchoconstriction. Of note, NPY1R was shown to drive immune cell infiltration in the airways^71^.

### Conclusion

In summary, our data revealed a new class of vagal airway specific nociceptors that acquire an inflammatory gene signature during allergic inflammation or when stimulated with IL-13. *Npy1r* overexpression was induced by IL-13 in a JAK/STAT6-dependent manner. During allergic airway inflammation, sympathetic nerve fibers and M2 macrophages release NPY, which subsequently decrease nociceptor neurons’ activity (**SF. 9**) and impact the levels of allergic airway inflammation. Future work will reveal whether targeting vagal NPY1R constitutes a relevant therapeutic target to quell asthma-induced bronchoconstriction and cough.

---

**Supplementary Figure 1.** <u>Airway vagal nociceptor neurons have a unique transcriptome.</u>

**(A)** Jugular nodose complex neurons gating strategy. Small debris were eliminated (FSC/SSC), and the whole cells were identified (nucleus marker SYTO40). Populations of airway-innervating nociceptor neurons (Na_V_1.8^+^DiD^+^), visceral nociceptors (Na_V_1.8^+^DiD^-^) and Na_V_1.8^-^ cells were then separated. Lumbar DRG were used as gating controls since they do not innervate the airways **(A)**.

**(B)** The transcriptome of purified airway-innervating nociceptor neurons (Na_V_1.8^+^DiD^+^), visceral nociceptors (Na_V_1.8^+^DiD^-^) and glial cells (Na_V_1.8^-^DiD^-^) was analyzed by RNA sequencing and population segregation was confirmed using principal component analysis **(B)**.

**(C)** Naive 8-week-old male and female Na_V_1.8^cre^::tdTomato^fl/wt^ mice were injected intranasally with the retrograde tracer DiD’ (200 µM). Fourteen days later, the mice were euthanized and JNC, thoracic DRG, and TG ganglia isolated, dissociated, and imaged with a fluorescence microscope. DiD’ retrotracer was detected in JNC nociceptor neurons but virtually absent in other ganglia (**C**).

(**D-E**) Naive 8-week-old male and female C57BL6 mice were injected intranasally with the retrograde tracer DiD’ (200 µM). Fourteen days later, the mice were euthanized and their JNC ganglia isolated and dissociated. JNC neurons were cultured (16 h) and calcium responsiveness to noxious stimuli was assessed. While the average neuronal responsiveness to the TRPV1 agonist capsaicin (300 nM) was stable between the two groups (**D**), the calcium flux induced by the TRPA1 agonist JT010 (50 µM) was higher in airway-innervating nociceptor neurons (**E**).

*Data are shown as flow cytometry dot plot (**A**), principal component analysis (**B**), and mean ± S.E.M (**C–E**). N are as follows: **C**: n=4 culture dishes**, D**: n=116 airway-innervating neurons and 1307 visceral neurons, **E**: n=137 airway-innervating neurons and 1406 visceral neurons. P-values were determined by a two-sided unpaired Student’s t-test (**C**) and are indicated in the figure*.

**Supplementary Figure 2.** <u>JNC airway nociceptor neuron markers</u>

(**A-D**) UMAPs of *Slc17a6*^+^ (VGLUT2) JNC neurons from single-cell RNA sequencing showing expression of *Phox2b* (**A**), *Prdm12* (**B**), *Scn1a* (**C**), and *Scn10a* (**D**). *Phox2b* and *Prdm12* delineate the nodose and jugular neurons, while *Scn1a* and *Scn10a* delineate low-threshold sensory neuron and nociceptor neuron populations. The experimental details were defined in *Prescott et al.* and the bioinformatic analysis is described in the methods section (**E**).

(**E–J**) Violin plot showing expression of *Npy1r* (**E**), *Kcng1* (**F**), *Trpa1* (**G**), *Il6* (**H**), *Il13ra1* (**I**), *Stat6* (**J**) in neuronal cells for each cluster identified by single-cell sequencing. All these genes are co-expressed in the airway-specific nociceptor neuron cluster NN8. The experimental details were defined in *Prescott et al.* and the bioinformatic analysis is described in the methods section.

*Data are shown as UMAPs with log-normalized expression as a feature (**A-D**) or as a violin plot of the log-normalized expression (**E-J**)*.

**Supplementary Figure 3.** <u>Lung projecting neurons single cell RNA sequencing.</u>

(**A-B**) UMAPs of barcode positive JNC neurons cluster from single-cell RNA projection-sequencing^7^ (**A**) and lung barcode expression (**B**). 4 clusters innervate the lung, 2 of them being lung specific, the two others also innervating stomach and colon (**B**). The experimental details were defined in Zhao *et al*^7^. and the bioinformatic analysis is described in the methods section.

(**C-H**) Violin plot showing expression of *Npy1r* (**C**), *Kcng1* (**D**), *Trpa1* (**E**), *Il6* (**F**), *Il13ra1* (**G**), *Scn10a* (**H**) in neuronal cells for each cluster identified by single-cell sequencing.

*Data are shown as UMAPs with log-normalized expression as a feature (**A-B**) or as a violin plot of the log-normalized expression (**C-H**)*.

**Supplementary figure 4.** <u>Single cell RNA seq reveals immune cells heterogeneity upon AAI.</u>

(**A**) 8-week-old male and female nociceptor neuron reporter (Na_V_1.8^cre^::tdTomato^fl/wt^) mice underwent the ovalbumin mouse models of asthma. Allergic inflammation was induced in mice by an initial sensitization to ovalbumin (OVA) (i.p. day 0 and 7) followed by inhaled OVA challenges (days 14–17). On days 2, 3, and 4, mice were injected intranasally with the retrograde tracer DiD’ (200 µM). One day after the last allergen challenge, the mice were euthanized and their JNCs isolated and dissociated, and airway-innervating nociceptor neurons (Na_V_1.8^+^DiD^+^) were purified by flow cytometry (**A**).The number of JNC airway-innervating nociceptor neurons is similar between naïve and OVA-exposed mice (tdTomato^+^DiD^+^) (**A**).

**(B-C)** 8-week-old male and female C57BL6 mice underwent the ovalbumin model of asthma. Lungs were harvested, immune cells sorted by flow cytometry before analysis by scRNA-seq. 21 different subtypes of immune cells were found in the airways **(E)**. Various specific markers allow to identify those cell types including B cells (*Cd19+*), T cells (*Cd3e+*), Granulocytes (*S100ab+*), NK cells (*Klrk1+*), Basophils (*Cd200r3+*), and Antigen Presenting Cells (APC, which include Macrophages and Dendritic cells, with varying expression of *Cd68*, *Itgax* and *Lgals3*). **(F)**.

*Data are shown as mean ± S.E.M (**A**), Umap **(B),** or Umap displaying Seurat normalized gene expression for the indicated genes (**C**). N are as follows: **A**: n=3 biological replicates (4 mice per sample), **B-C**: n=1-3 biological replicates (2 mice per sample). P-values were determined by a two-sided unpaired Student’s t-test (**A**)*.

**Supplementary figure 5.** <u>IL-13 reprogram airway nociceptor neurons through IL4RII.</u>

(**A**) Naive 8-week-old male and female nociceptor neurons reporter (Na_V_1.8^cre^::tdTomato^fl/wt^) mice were injected intranasally with the retrograde tracer DID’ (200 µM). Fourteen days later, the mice were euthanized and their JNC ganglia isolated and dissociated. Airway-innervating nociceptor neurons (Na_V_1.8^+^DiD^+^), visceral nociceptors (Na_V_1.8^+^DiD^-^) and Na_V_1.8^-^ cells were purified by flow cytometry and RNA sequenced. Both IL4RII subunits, *Il4ra* and *Il13ra1*, were detected in JNC nociceptors. Other IL-13 and IL-4 receptors are not detected. *IL13ra1* transcript expression was higher in nociceptor neurons when compared to levels measured in Na_V_1.8^-^ cells (**A**).

(**B**) Naive 8-week-old male and female C57Bl6 mice were euthanized and their JNC and DRG ganglia were isolated and cultured (16 h). The neurons were then loaded with the calcium indicator Fura-2AM (5 µM) and their responsiveness to IL-13 (100 ng/mL) was assessed using calcium microscopy. DRG and JNC neurons show limited calcium response when exposed to IL-13 compared to its vehicle **(B)**.

(**C**) Naive 8-week-old male and female C57BL6 mice DRG neurons were isolated and cultured in the presence of IL-13 (100 ng/mL; 24 h) or its vehicle. Transcript levels were assessed by qPCR. In comparison to the vehicle, IL-13 increased DRG neurons’ expression of *Npy1r* (**C**).

*Data are shown as mean ± S.E.M (**A-B**), or as box (25th-75th percentile) and whisker (min-to-max) plots (**C**). N are as follows: **A**: n=3 biological replicates (4 mice per sample), **B**: n=11 culture dishes for JNC and 3 culture dishes for DRG, **C**: n=4 cultures from different mice per group. P-values were determined by one-way ANOVA with post hoc Tukey’s (**A**); or a two-sided unpaired Student’s t-test (**B-C**). P-values are shown in the figure*.

**Supplementary Figure 6.** <u>NPY is expressed in M2 macrophages.</u>

(**A-B**) Eight-week-old female mice were subjected to the ovalbumin-induced asthma model. Mice were initially sensitized to ovalbumin (OVA) intraperitoneally (i.p.) on days 0 and 7, followed by inhaled OVA challenges from days 14 to 17. On day 18, lungs were harvested and CD45^+^ cells were sorted by flow cytometry. Their transcriptomes were analyzed by single-cell RNA sequencing. Related to Figure 6, showing that NPY is expressed in the cluster of myeloid cells including Macrophages and Denditic cells, we show here that the NPY positive population also express M2 macrophages markers (*Fcgr1, Ccr5, Cd63, Mrc1, Msr1* and *Ccl24*).

**Supplementary Figure 7.** <u>NPY1R blunts JNC nociceptors excitability.</u>

**(A-C)** 8-week-old male and female NPY1R reporter (NPY1R^cre^::tdTomato^fl/wt^) mice were sacrificed and their JNC neurons harvested and cultured (16 hours). Whole cell patch clamp electrophysiology was performed on the NPY1R^+^ nociceptor neurons. A current clamp was applied while the neurons’ membrane potential was recorded before and after exposing (10 min) the cell to Leu^31^Pro^34^NPY (250 nM) or its vehicle. The number of action potentials for each neuron was normalized by the maximum number observed at baseline (**B-C**. While the vehicle had little to no effect on neuronal excitability (**A**, **B**), Leu^31^Pro^34^NPY reduced the number of action potentials in response to current stimulation in NPY1R^+^ (tdTomato) neurons **(A, C).**

*Data are shown as traces of membrane potential for individual neurons (**A**), mean ± S.E.M **(B, C)**, or box (25th–75th percentile) and whisker (min-to-max) plots **(D-E)**. N are as follows: **B, D**: n=10 vehicle treated neurons**, C, D:** n=9 Leu^31^Pro^34^NPY treated neurons, **E**: n=19 culture wells. P-values were determined by two-way ANOVA (**B, C**) or two-sided unpaired Student’s t-test **(D-E)**. P-values are shown in the figure*.

**Supplementary Figure 8.** <u>NPY1R does not affect immune cell infiltration but changes T cells profile.</u>

(**A-B**) Eight-week-old female nociceptor neuron NPY1R conditional knockout (Na_V_1.8^cre^::NPY1R^fl/fl^) and littermate control mice (Na_V_1.8^wt^::NPY1R^fl/fl^) were subjected to the ovalbumin-induced asthma model. Mice were initially sensitized to ovalbumin (OVA) intraperitoneally (i.p.) on days 0 and 7, followed by inhaled OVA challenges from days 14 to 17. One day after the last allergen challenge, bronchoalveolar lavage fluid (BALF) was collected and immunophenotyped using flow cytometry. OVA-exposed mice demonstrated significant airway inflammation with increased infiltration of leukocytes (CD45^+^), including eosinophils (CD45^+^CD11C^low^SiglecF^Hi^), neutrophils (CD45^pos^SiglecF^low^CD11B^high^LY6G^high^), and CD4 T cells (CD45^pos^SiglecF^low^CD11B^low^CD4^high^), while the number of alveolar macrophages (CD45^+^CD11C^high^SiglecF^Hi^) decreased. The conditional knockout of NPY1R did not affect BALF cell numbers (**A-B**).

(**C-H**) Eight-week-old female C57BL/6 mice, either pretreated with Resiniferatoxin or untreated, together with nociceptor neuron NPY1R conditional knockout mice (Na_V_1.8^cre^::NPY1R^fl/fl^) and their littermate controls (Na_V_1.8^wt^::NPY1R^fl/fl^), were subjected to the ovalbumin-induced asthma model. On day 18, lungs were harvested and CD45^+^ cells were sorted by flow cytometry. Their transcriptomes were analyzed by single-cell RNA sequencing. T-cells were subclustered, and various markers were used to classify the different subpopulations. UMAPs from each biological replicate of OVA-exposed Na_V_1.8^wt^::NPY1R^fl/fl^ (**D**), Na_V_1.8^cre^::NPY1R^fl/fl^ (**E**), C57BL6 mice treated with vehicle (**F**), C57BL6 mice treated with RTX (**G**), and naive C57BL6 (**H**) were produced (**C-H**).

Data are presented as mean ± S.E.M (**A-D**). Sample sizes are as follows: **A-B**: n=17 mice per group; **D-E**: n=3 samples (2 mice per sample), **F-G**: n=2 samples (2 mice per sample), **H**: n=1 sample (2 mice per sample)P-values were determined one-way ANOVA with post hoc Tukey’s test (A, B). P-values are indicated in the figure.

**Supplementary figure 9.**
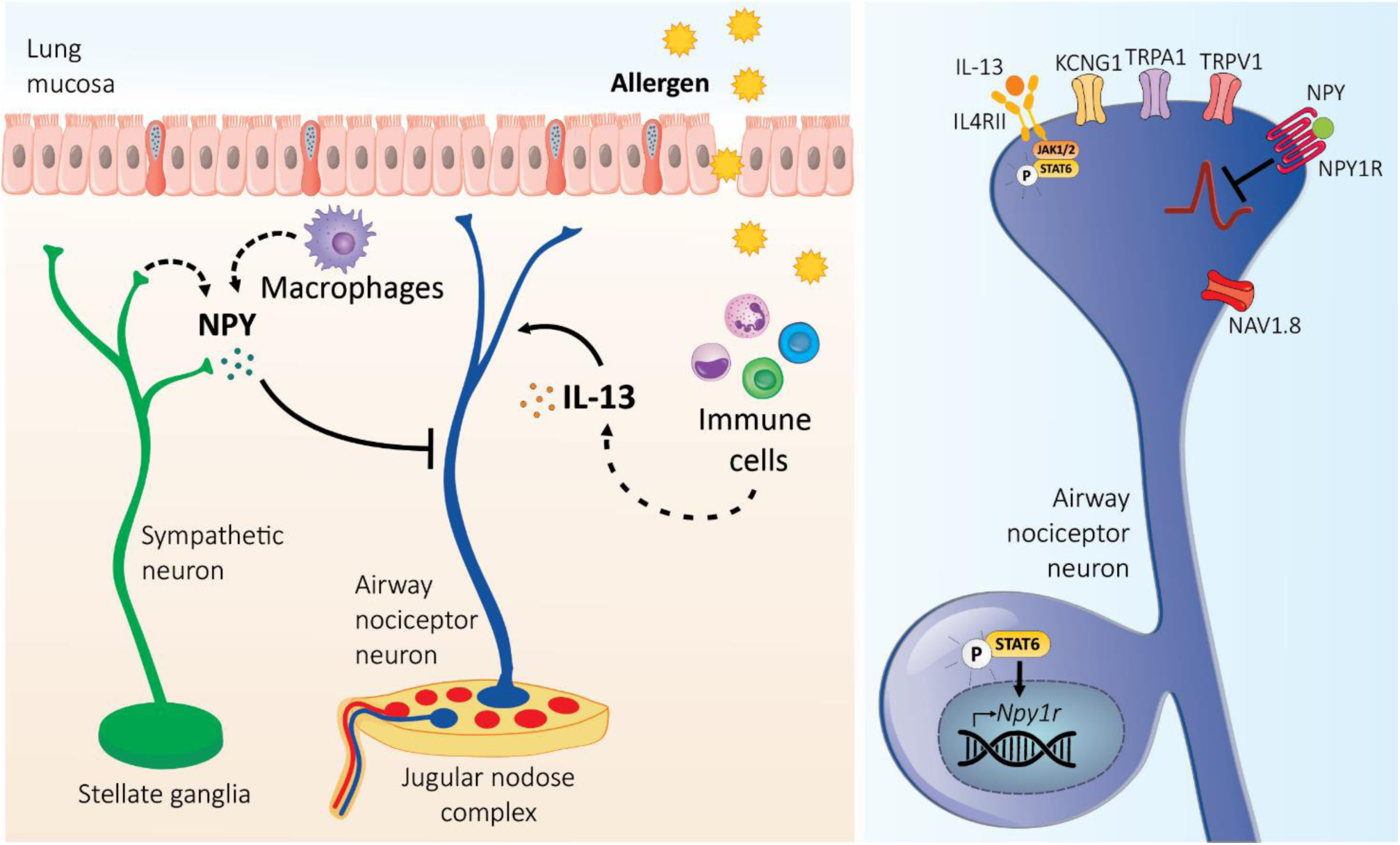
IL-13 promotes sensory-sympathetic neurons crosstalk in asthma. When allergens are present in the airways, the immune and stromal cells in the area are activated and release inflammatory cytokines such as IL-13. These cytokines are then detected by nociceptor neurons, which leads to specific changes in gene expression. For example, IL-13 signaling through its interaction with IL4RII leads to increased expression of Npy1r via phosphorylation of STAT6. Via the action of neuropeptide Y (NPY) released by sympathetic neurons and macrophages, this cascade ultimately decreased the sensitivity of NPY1R-expressing nociceptor neurons through reduced intracellular levels of cAMP.

**Supplementary Table 1.** <u>Airway neuronal subtypes</u>. Single cell sequencing data of JNC cells from Prescott et al. were reanalyzed using Seurat. Neuronal cells were selected based on *Slc17a6* (Vglut2) expression. A total of 21 neuronal populations were identified. *Phox2b* and *Prdm12* were used to identify nodose and jugular groups, while nociceptor neurons and low-threshold sensory neurons were defined based on their expression of *Scn10a* and *Scn1a*. The markers identified for each cluster were compared to airway-innervating neurons sequencing by GSEA. A positive normalized enrichment score indicates preferential innervation of the airways for a given neuronal population. Detailed classification of neurons preferentially innervating the airways is displayed in a tab. Other tabs display the average expression of all genes for all neuronal clusters and DESeq2 analysis comparing airway and visceral nociceptor neurons. Another tab shows the average gene expression in clusters identified in reanalysis of projection-seq data from Zhao et al.

**Supplementary Table 2.** <u>Single cell sequencing of lung immune cells of naïve and AAI mice.</u> 8-week-old female C57BL6 mice underwent the ovalbumin mouse models of asthma. Allergic inflammation was induced in mice by an initial sensitization to ovalbumin (OVA) (i.p. days 0 and 7) followed by inhaled OVA challenges (days 14–17). One day after the last allergen challenge, the lung was harvested, CD45+ cells sorted by flow cytometry before analysis by single cell RNA sequencing. The tables displays the differentially expressed genes induced by AAI, determined using Seurat.

**Supplementary Table 3.** <u>DESeq2 analysis of JNC cell populations in AAI and Naive conditions</u>. 8-week-old male and female nociceptor neuron reporter (Nav1.8^cre^::tdTomato^fl/wt^) mice underwent the ovalbumin mouse models of asthma. Allergic inflammation was induced in mice by an initial sensitization to ovalbumin (OVA) (i.p. days 0 and 7) followed by inhaled OVA challenges (days 14–17). On day 2, 3, and 4, mice were injected intranasally with the retrograde tracer DiD’ (200 μM). One day after the last allergen challenge, the JNC neurons were harvested for flow cytometry purification and RNA sequencing. DESeq2 pairwise comparison of airway-innervating (Na_V_1.8^+^DiD^+^) naive nociceptor neurons versus AAI nociceptor neurons counterparts shows 92 differentially expressed genes. A literature review for each gene allowed a classification of their expected function ^67,108–158^. Other tabs display DESeq2 analysis of visceral nociceptors (Na_V_1.8^+^DiD^+^) and Na_V_1.8^-^ cells.

**Supplementary Table 4.** <u>Cytokines reprograms nociceptor transcriptome.</u> 8-week-old male and female nociceptor neuron reporter (Na_V_1.8^cre^::tdTomato^fl/wt^) mice JNC neurons were cultured (24 h) with IL-13 (100 ng/mL), IL-1β (100 ng/mL), BDNF (50 ng/mL), TNF-α (100 ng/mL) or vehicle, before nociceptor purification by flow cytometry and RNA-sequencing. Data from Cobos and colleagues ^53^ was also reanalyzed by DESeq2 and shows various differentially expressed genes induced in whole DRG three days after spared nerve injury. The different tabs show the DESeq2 analysis for each of these conditions.

**Supplementary Table 5.** <u>DEGs in IL-13 exposed nociceptors</u>. 8-week-old male and female nociceptor neuron reporter (Na_V_1.8^cre^::tdTomato^fl/wt^) mice JNC neurons were cultured (24 h) with IL-13 (100 ng/mL) or vehicle. The nociceptor neurons were then purified by flow cytometry and changes to their transcriptome analyzed by RNA sequencing. The tables display the results of the DESeq2 analysis comparing the vehicle and IL-13-exposed conditions. DEGs are showed in a separate tab. A literature review for each gene allowed a classification of their expected function^67,159–181^.

**Supplementary Table 6.** <u>FPKM normalized sequencing data</u>. Fragment per kilobase per million (FPKM) normalized values for all JNC sequencing data produced in this study (airway nociceptors, visceral nociceptors, Na_V_1.8^-^ cells in naive and allergic airway inflammation conditions, cultured nociceptors exposed to vehicle, cytokines, or neurotrophins).

**Supplementary Table 7.** <u>OVA-exposed Na_V_1.8^cre^::NPY1R^fl/fl^ mice show differential gene expression in lung immune cells compared to littermate control</u>. 8-week-old female mice with NPY1R conditional knock-out in nociceptors (Na_V_1.8^cre^::NPY1R^fl/fl^) and littermate controls (Na_V_1.8^wt^::NPY1R^fl/fl^) underwent the ovalbumin mouse models of asthma. Allergic inflammation was induced in mice (2 mice per samples, 3 samples per group) by an initial sensitization to ovalbumin (OVA) (i.p. days 0 and 7) followed by inhaled OVA challenges (days 14–17). One day after the last allergen challenge, the lung was harvested, CD45+ cells sorted by flow cytometry before analysis by single cell RNA sequencing. To identify robust gene expression changes, a pseudo-bulk approach was used. Cell types were identified using common immune markers. Shown are gene expression averaged for each cell type in each sample identified between OVA-exposed Na_V_1.8^wt^::NPY1R^fl/fl^ and Na_V_1.8^wt^::NPY1R^fl/fl^ mice.

**Supplementary Table 8.** <u>OVA-exposed Na_V_1.8^cre^::NPY1R^fl/fl^ mice show differential gene expression in lung immune cells compared to littermate control</u>. 8-week-old female mice with NPY1R conditional knock-out in nociceptors (Na_V_1.8^cre^::NPY1R^fl/fl^) and littermate controls (Na_V_1.8^wt^::NPY1R^fl/fl^) underwent the ovalbumin mouse models of asthma. Allergic inflammation was induced in mice (2 mice per samples, 3 samples per group) by an initial sensitization to ovalbumin (OVA) (i.p. days 0 and 7) followed by inhaled OVA challenges (days 14–17). One day after the last allergen challenge, the lung was harvested, CD45+ cells sorted by flow cytometry before analysis by single cell RNA sequencing. To identify robust gene expression changes, a pseudo-bulk approach was used. Cell types were identified using common immune markers. Shown are differentially expressed genes identified between OVA-exposed Na_V_1.8^wt^::NPY1R^fl/fl^ and Na_V_1.8^wt^::NPY1R^fl/fl^ mice.

**Supplementary Table 9.** <u>OVA-exposed Na_V_1.8^cre^::NPY1R^fl/fl^ mice show differential gene set enrichment in lung immune cells compared to littermate control.</u> 8-week-old female mice with NPY1R conditional knock-out in nociceptors (Na_V_1.8^cre^::NPY1R^fl/fl^) and littermate controls (Na_V_1.8^wt^::NPY1R^fl/fl^) underwent the ovalbumin mouse models of asthma. Allergic inflammation was induced in mice (2 mice per samples, 3 samples per group) by an initial sensitization to ovalbumin (OVA) (i.p. days 0 and 7) followed by inhaled OVA challenges (days 14–17). One day after the last allergen challenge, the lung was harvested, CD45+ cells sorted by flow cytometry before analysis by single cell RNA sequencing. To identify robust gene expression changes, a pseudo-bulk approach was used. Cell types were identified using common immune markers. Shown are GO gene set enrichment for each cell type identified between OVA-exposed Na_V_1.8^wt^::NPY1R^fl/fl^ and Na_V_1.8^wt^::NPY1R^fl/fl^ mice.

**Supplementary Table 10.** <u>T cell subpopulations analysis</u>. T cells were subsetted from the CD45+ single cell RNA sequencing data. High resolution clustering revealed several clusters of T cells and innate lymphoid cells, including CD4, CD8, iT-cells (immature T cells), ILCs, NKT, and γδT cells. The table displays average expression in each of those lymphocytes cell subtypes, in each sample, as well as the top markers of each cluster.

## METHODS

### Animals

Mice were housed in standard environmental conditions (12h light/dark cycle; 23°C; food and water ad libitum) at facilities accredited by CCPA. Parental strain C57BL6 (Jax, #000664), tdTomato^fl/fl^ (Jax, #007914), Trpv1^cre/cre^ (Jax, #017769), Npy1r^cre/cre^ (Jax, #030544) were purchased from Jackson Laboratory. Parental strain Nav1.8^cre/cre^ mice were generously supplied by Prof. John Wood (UCL). Parental strain NPY1R^fl/fl^ mice were generously supplied by Prof. Herbert Herzog (Gavan Institute of Medical Research). Male and female mice were bred in-house and used between 6 and 12 weeks of age. Cross breeding was used to generate the following genotypes: Na_V_1.8^cre^::tdTomato^fl/wt^, NPY1R^cre^::tdTomato^fl/wt^, TRPV1^cre^::tdTomato^fl/wt^, Na_V_1.8^cre^::NPY1R^fl/fl^, and Na_V_1.8^wt^::NPY1R^fl/fl^ (littermate control).

### Ovalbumin model of allergic airway inflammation

On days 0 and 7, mice were sensitized by a 200 µL i.p. injections of a solution containing 1 mg/mL ovalbumin (Sigma, #A5503) and 5 mg/ml aluminum hydroxide (Sigma, # 239186). On days 15–17, mice were lightly anesthtetized (isoflurane 2.5 %, CDMV #108737) and instilled daily with 50 μg/50 µL OVA intranasally. Mice were sacrificed on day 18 unless otherwise indicated.

### House dust mite model of allergic airway inflammation

Lightly anesthetized (isoflurane 2.5 %, CDMV #108737) mice were challenged (20 μg/50 μL, intranasal) on day 1–5 and 8–10 with house dust mites (CiteQ Biologics, #15J01) and sacrificed on day 11.

### Airway neurons’ retrograde tracing

Mice were injected (50 μL, 200 μM in PBS, 1% DMSO; intranasally) with DiD’ (Thermofisher, #D7757) for 3 consecutive days (D2-D4 of OVA protocol). The mice were sacrificed two weeks after the last injection to allow the tracer to reach the JNC.

### AS1517499

Ovalbumin was mixed with AS1517499 (150 μg / 50 μL) or vehicle (DMSO), for intranasal co-instillation at each challenge of the OVA protocol (day 15-16-17). Naïve mice received the vehicle (DMSO) without Ovalbumin.

### 6-OHDA

6-Hydroxydopamine was injected at 12.5 mg/mL then 37.5 mg/mL (200ul I.P. in 0.1% ascorbic acid) respectively 5 and 3 days before the first OVA sensitization, then again 5 and 3 days before the first OVA intranasal challenge.

### Neuron culture

Mice were sacrificed and JNC or DRG were dissected into an ice-cold HEPES buffered DMEM medium (Thermofisher, #12430062). JNC ganglia from 2-10 mice were pooled in the same tube. The cells were transferred to a HEPES buffered DMEM medium completed with 1 mg/mL collagenase IV (Sigma, #C5138) + 2.4 U/mL dispase II (Sigma, #04942078001) and incubated for 70 minutes at 37 °C. Ganglia were triturated with glass Pasteur pipettes of decreasing size in DMEM medium, then centrifuged (200 g) over a 15% BSA gradient in PBS to eliminate debris. The neurons were plated on laminin (Sigma, #L2020) coated cell culture dishes. The cells were cultured at 37° with Neurobasal-A medium (Gibco, #21103-049), supplemented with 2% B27 (Thermofisher, #17504044) 0.01 mM AraC (Sigma, #C6645), 200 nM L-Glutamine (VWR, # 02-0131), 100 U/mL penicillin and 100 μg/mL streptomycin (Corning, #30002CI) without neurotrophin unless otherwise indicated. Culture densities and durations are described for each application.

### Cyclic AMP

JNC neurons were cultured (1.5×10^3^) in 96 well plates (VWR #10062-900) for 16 hours. The media was removed, and neurons exposed to Leu^31^Pro^34^NPY (250nM Tocris #1176) or vehicle (PBS containing cAMP phosphodiesterase inhibitor (100 μM Ro-20-1724; Sigma, #557502 and 500 μM 3-Isobutyl-1-methylxanthine; Sigma, #I5879) for 30 minutes. Cells were lysed for enzymatic cAMP measurement assay using a commercial kit and following manufacturer’s instructions (Promega, #V1501).

### Calcium microscopy

C57BL6 or NPY1R^cre^::tdTomato^fl/wt^ DRG or JNC neurons were plated (2×10^3^) on laminin-coated glass-bottom dishes (35 mm; ibidi, #81218) and cultured overnight. The cells were then loaded with Fura-2-AM (5 μM, 37 °C, Biovision, #125280) for 45min, washed with Standard Extracellular Solution (145 mM NaCl, 5 mM KCl, 2 mM CaCl_2_, 1 mM MgCl_2_, 10 mM glucose, 10 mM HEPES, pH 7.5), and imaged at room temperature. Capsaicin 50 nM, 100 nM and 300 nM (Tocris, #0462) or JT010 10 μM and 50 μM (Sigma, #SML1672), or IL-13 (100 ng/mL, Biolegend #575904) were prepared in SES and were flowed (30 seconds, 1 minute and 1 minute, respectively) directly onto neurons using perfusion barrels followed by buffer washout (5 minutes). 40 mM KCl solution was then flowed on the cells for 20 seconds. To test NPY1R effect on neuron sensitivity, Leu^31^Pro^34^NPY (250 nM; Tocris, #1176) or vehicle was perfused for 5 minutes before the other compounds. A single field of view was acquired per dish. Cells were illuminated by a UV light source (Cool LED, pE-340) with alternating 340 nm and 380 nm excitation, and a camera (Photometrics Prime 95B 25 mm) captured fluorescence emission (515/45 nm) with a 20X objective on a Ti2 microscope (Nikon). Regions of interest (ROI) were manually drawn and 340/380 fluorescence ratios were exported. Microsoft Excel was used for further analysis (Microsoft, USA). Neurons were considered responsive to a compound if the fluorescence ratio increased by at least 10% within 1 minute after injection.

### Electrophysiology

JNC neurons from NPY1R^cre^::tdTomato^fl/wt^ mice were plated (2×10^3^) onto Poly-D-lysin and laminin-coated glass-bottom 35 mm dishes (Ibidi #81218) and cultured in supplemented neurobasal medium. TdTomato-positive JNC neurons were identified using a Nikon Eclipse Ti microscope. Whole-cell voltage and current-clamp recordings were performed with an EPC 800 patch clamp amplifier (HEKA) and were filtered at 10 kHz with the internal Bessel filter and digitized using an Axon Instruments Digidata 1440A digitizer at 20 kHz. The neurons were placed in standard external solution (163 mM NaCl, 2.5 mM CaCl_2_, 2 mM MgCl_2_, 20 mM glucose, and 10 mM HEPES, pH = 7.4). Recording pipettes were pulled from soda lime glass pipettes (outer diameter 1.5 mm, Kimble® Chase, #41A2502,) using a P-97 microelectrode puller (Sutter Instrument). Pipettes with resistances of 3–5 MΩ were used for recordings. Pipettes were filled with internal solution (133 mM potassium gluconate, 6 mM NaCl, 1 mM CaCl_2_, 0.7 mM MgCl_2_, 10 mM HEPES, and 10 mM EGTA, pH = 7.2, final K_in_^+^ concentration –163mM).

For voltage clamp experiments, the neurons were clamped at –60mV. Neuronal currents were recorded before and after drug exposure from a series of depolarization steps ranging from –120 to +70mV in 10 mV intervals. For the current clamp AP protocol, neurons were injected with a series of 35 pA current steps (1 s duration) from 0 to 630 pA. These voltage and current clamp experiments were performed before and 10 mins post-addition with a pipette of external solution (vehicle) or Leu^31^Pro^34^NPY (250 nM; Tocris, #1176). Current amplitudes and APs were quantified using Clampfit 10.7 (Molecular Devices). Since baseline activity varied among the JNC neurons, the activity of the neurons after application of vehicle or Leu^31^Pro^34^NPY was normalized to baseline activity of the same cell.

### Bronchoalveolar lavage fluid (BALF)

Mice were anesthetized by intraperitoneal injection of urethane (250 μL i.p., 20%) and a 20G sterile catheter inserted longitudinally into the trachea. 500 μL followed by 1 mL of ice-cold buffer (PBS, 2% FBS, 1 mM EDTA) containing protease inhibitors (Sigma, #P1860) was injected into the lung, harvested, and stored on ice. BALF underwent a 400g centrifugation (5 min; 4 °C), supernatant of the first flush was harvested and frozen (−80°C) for subsequent ELISA experiments. Cells were resuspended for cell count and flow cytometry analysis.

### BALF immunophenotyping

Cells were resuspended in red blood cells lysis buffer (Gibco, #A10492-01) for 1 minute then washed with PBS. Cells were then stained with Zombi Aqua (Biolegend, #423102) for 10 minutes, before staining in a flow cytometry buffer (PBS, 2% FBS, 1 mM EDTA) supplemented with 1% Rat Serum. Staining antibodies included anti-CD45–BV421 (1:400; BioLegend, #103134), anti-Siglec-F–PE (1:400; Thermofisher, #12-1702-82), anti-CD11b-APC–Cy7 (1:400, BioLegend, #101262), anti-CD11c–FITC (1:400, BioLegend, #117306), anti-Ly6G–APC (1:400, BioLegend, #127614), anti-Ly6C–PE-Cy7 (1:400, BioLegend #128018), and anti-CD4-PerCP–Cy5.5 (1:400, BioLegend, #116012) for 30 minutes in the dark at 4 °C. 1×10^4^ counting beads (Biolegend, #424902) were added for absolute quantification before data acquisition on a FACS Canto II (BD Biosciences). Leukocytes were gated as CD45^pos^, eosinophils as CD45^pos^SiglecF^high^CD11C^low^, alveolar macrophages as CD45^pos^SiglecF^high^CD11C^high^, neutrophils as CD45^pos^SiglecF^low^CD11B^high^LY6G^high^, and CD4 T cells as CD45^pos^SiglecF^low^CD11B^low^CD4^high^.

### Flow cytometry sorting of sensory neurons

Na_V_1.8^cre^::tdTomato^fl/wt^ mice were sacrificed and JNC or DRG were dissected into ice-cold HEPES buffered DMEM medium (Thermofisher, #12430062). For RNA sequencing of airway nociceptor neurons, JNC ganglia were pooled from 4 mice (2 males and 2 females) for each biological replicate. Ganglia were transferred into a HEPES buffered DMEM medium completed with 1 mg/mL collagenase IV (Sigma, #C5138) + 2.4 U/mL dispase II (Sigma, #04942078001) and incubated for 70 minutes at 37°C, then washed in DMEM medium before trituration with glass Pasteur pipettes of decreasing size. Cells were then centrifuged (200g) over a 15% BSA gradient in PBS to eliminate debris. Nucleus were stained with SYTO40 (10 μM, 5 minutes RT; Thermofisher, #S11351) to differentiate cells from axonal debris, washed with PBS, then cells were resuspended in sterile flow cytometry buffer (PBS, 2% FBS, 1 mM EDTA) and filtered (70 µm; VWR, #10204-924).

For DiD’-injected mice, JNC neurons were sorted directly into Trizol (500 μL; Invitrogen, #15596026) and stored at −80°C for subsequent RNA extraction. Airway nociceptors were gated as Syto40^hi^tdTomato^hi^DiD^pos^, visceral nociceptors as Syto40^hi^tdTomato^hi^DiD^lo^, and glial/stromal Na_V_1.8^-^ cells as Syto40^hi^tdTomato^lo^. DiD’-injected Na_V_1.8^cre^::tdTomato^fl/wt^ mice lumbar DRG neurons and naive Na_V_1.8^cre^::tdTomato^fl/wt^ mice JNC neurons were used as gating controls for DiD’. JNC neurons from C57BL6 mice were used as gating controls for tdTomato.

### BALF ELISA

Cell free BALF supernatant and serum were used for ELISA quantification after storage at −80°. IL-13 (Thermofisher, #88-7137-88) and NPY (Cusabio, #CSB-E08170M) were measured using commercial kits following manufacturer’s instructions.

### Nociceptor neuron ELISA

C57BL6 mice JNC neurons were cultured (1.5×10^4^) in 96 well plates with vehicle or IL-1β (100 ng/mL). After 36 hours, supernatant was harvested, centrifuged (500g) to remove cellular debris, and IL-6 secretion analyzed using a commercial ELISA kit following the manufacturer’s instructions (Biolegend, #431304).

### Whole-lung RNA extraction

Whole lungs were minced with a razor blade, and about a quarter of the minced lung was mixed with 500uL Trizol (Thermofisher #15596018) and stored at −80° for subsequent RNA extraction. RNA was separated from protein and DNA by mixing 500 μL of sample in Trizol with 100 μL chloroform before ultracentrifugation (15 min, 16000g, 4°). The upper phase was mixed with a half volume of 100% isopropanol, transferred to a purification column, and the RNA was then purified using the kit E.Z.N.A. Total RNA Kit I (VWR, #CA101319) following manufacturer’s instructions.

### JNC Neurons RNA extraction

RNA was separated from protein and DNA by mixing 500 μL sample in Trizol with 100 μL chloroform before ultracentrifugation (15 min, 16000g, 4°). The upper phase was mixed with a half-volume of 100° isopropanol, transferred to a purification column, and the RNA purified using the kit PureLink RNA Micro Scale (Thermofisher, #12183016) following manufacturer’s instructions.

### Screening of cytokine neuromodulatory capacity

JNC neurons were plated (5×10^3^; 200 μL) on laminin-coated 96 well plates and exposed to cytokines (100 ng/mL; IL-1β, IL-4, IL-5, IL-6, IL-10, IL-13, IL-17, IL-31, IL-33, SCF, TGF-β, TSLP or TNF-α), lipid mediators (200 ng/mL; leukotriene C4), neurotrophins (50 ng/mL; nerve growth factor or brain-derived neurotrophic factor), IgE-OVA (10 μg/mL) or NPY (100 nM). To investigate IL-13 intracellular signaling, JNC neurons were incubated with IL-13 (100 ng/mL, Biolegend #575904) for 24 hours in presence or not of the STAT6 inhibitor AS1517499 (1 μM MCE #HY-100614) or the JAK1/2 inhibitor Ruxolitinib (10 μM, MCE #HY-50856). Biological replicates were made using different mice preparations for each replicate. After 24 hours of culture, the medium was removed, and the cells harvested in 500 μL Trizol (Thermofisher, #15596018). Samples were then used for RNA extraction and RT-qPCR.

### RT-qPCR

RNA was reverse transcribed using the SuperScript VILO Master Mix (Thermofisher, #11755250). The cDNA was then subjected to two-step thermocycling using PowerUp qPCR SYBR Green Mix (Invitrogen, #A25742) and data collection was then performed on a Mic qPCR machine (Bio Molecular Systems). 1–3 ng cDNA was used for qPCR from neuron preparations, and 80–100 ng for qPCR from whole lung RNA. Expression levels were normalized using the ΔΔCt method with *Actb* as the reference gene.

The primers used were : *Actb* Forward: TGTCGAGTCGCGTCCACC; *Actb* Reverse: TATCGTCATCCATGGCGAACTGG; *Trpa1* Forward: TCCAAATTTTCCAACAGAAAAGGA; *Trpa1* Reverse: CGCTATTGCTCCACATTGCC; *Npy1r* Forward: CTCGTCCCGCTTCAACAGAG; *Npy1r* Reverse: TCAAAACGGATCAAATCTTCAGCA; *Il6* Forward: GGATACCACTCCCAACAGACC; *Il6* Reverse: AATTGCCATTGCACAACTCTTTTC; *Bdnf* Forward: CAGGTTCGAGAGGTCTGACG; *Bdnf* Reverse: AAGTGTACAAGTCCGCGTCC; *Sting1* Forward: CTGCCGGACACTTGAGGAAA; *Sting1* Reverse: CCGTCTGTGGGTTCTTGGTAG; *Npy* Forward: CGCTCTGCGACACTACATCA; *Npy* Reverse: AGGGTCTTCAAGCCTTGTTCT; *Il13* Forward: CCAGACTCCCCTGTGCAAC; *Il13* Reverse: GGCTACACAGAACCCGCC; *Muc5ac* Forward: AAGATCAACTGTCCGCAGGG; *Muc5ac* Reverse: GGTGCACCGTACATTTCTGC; *Muc5b* Forward: AAGAGCACAGAGTGTCAGGC; *Muc5b* Reverse: GTGTGTGGCAGCTTTTGAGG.

### Western Blot

C57BL6 mice DRG or JNC neurons (1.5×10^4^; 200 μL) were plated onto 35mm well plates and cultured overnight. Neurons were exposed to IL-13 or vehicle for 60 minutes (37°), then washed with PBS and lysed in RIPA buffer (Sigma, #20-188) supplemented with phosphatase inhibitor (1/100, Sigma #P0044) and protease inhibitor (1/200, Sigma, #P1860) for 20 minutes at room temperature. Protein concentrations were measured by BCA (Thermofisher, #23227) and equilibrated with a supplemented RIPA buffer. Samples were then mixed 1:1 with loading buffer (0.1M Tris pH 6.8; 4% SDS; 0.25% Bromophenol blue; 20% glycerol; 10% β-mercaptoethanol) and heated to denature proteins (100 °C, 5 minutes). 10 μg of protein were loaded and separated by SDS-PAGE electrophoresis with an acrylamide gel. Proteins were then transferred to a nitrocellulosis membrane, blocked with blocking buffer (TBS-T buffer containing 5% BSA), incubated with primary antibodies in blocking buffer (overnight, 4°), washed, and incubated with secondary antibodies in blocking buffer (1 h, RT). Membranes were then washed and revealed using SuperSignal West Dura Extended Duration Substrate (Thermofisher, #34075). Antibodies used included rabbit anti-pSTAT6 (1/1000, CST, #9361S), rabbit anti-STAT6 (1/1000, CST, #9362S) and HRP anti-rabbit (1/2000, CST, #7074S).

### Neurite growth assay

Na_V_1.8^cre^::tdTomato^fl/wt^ JNC neurons were plated (1.5×10^3^) on laminin-coated glass-bottom dishes (35 mm; ibidi, #81218) and cultured for 24 hours. Neurobasal media was supplemented or not with with IL-13 (100 ng/mL, Biolegend #575904), IL-1β (100 ng/mL, Biolegend #575104), TNF-α (100 ng/mL Biolegend # 575204), BDNF (50 ng/mL Peprotech #450-02) or vehicle. Pictures of the whole plating area were taken after 24 hours of culture using a 20X objective to collect tdTomato fluorescence (Excitation 554/23nm; Emission:609/54nm). For the analysis of neurite outgrowth, an in-house developed method was used. Using Nikon Elements software, a fluorescence threshold was used to define the tdTomato positive neurites and soma. The somas were then defined and excluded based on fluorescence intensity, size, and circularity. The total size of neurites was then divided by the number of somas for each culture dish.

### Immunofluorescence

C56BL6 and Na_V_1.8^cre^::tdTomato^fl/wt^ mice were anesthetized with urethane (20%, 250 μL, i.p.) and perfused with 10 mL PBS, then with 10 mL 4% PFA. Lungs, JNC, DRG and SCG were harvested and incubated in 4% PFA for 24 hours at 4°. Organs were then sequentially transferred in 10%, 20%, then 30% sucrose (24 hours each), before mounting and freezing (−80°) in OCT. Cryosections of 40 μm for lungs and 15 μm for ganglia were prepared. For neuron cultures, the medium was removed, and the cells fixed with 4% PFA for 15 minutes. Sections and cultures were incubated in blocking solution (1X PBS, 0.2% Triton X-100, 50 mg/mL BSA and 5% goat serum) for 3 hours at room temperature, then incubated with primary antibodies in staining solution (PBS, 0.2% Triton X-100 and 3% goat serum) for 48 hours at 4°. Primary antibodies used included rabbit anti-NPY (1/500, CST #11976S), guinea pig anti-PGP9.5 (1/1000, Sigma #AB5898), chicken anti-TH (1/500, Aves Laboratory #TYH-0020), rat anti-CD45 BV421 conjugated (1/300, Biolegend ##103134), rabbit anti-IL6 (1/300, CST #12912T). Cryosections were then washed (3 times, 15 minutes with shaking) with PBS containing 0.2% Triton X-100. Secondary antibodies were incubated in staining solution for 16 hours at 4°. Secondary antibodies included AF488 goat anti-guinea pig (1/1000, Thermofisher #A11073), AF546 goat anti-guinea pig (1/1000, Thermofisher #A-11074) AF647 goat anti-rabbit (1/1000, Thermofisher #A21245), CF488 goat anti-chicken (1/1000, Sigma #SAB4600039), AF405 goat anti-chicken (1/1000, Thermofisher #A48260). Ti2 microscope (Nikon) equipped with a camera (Photometrics Prime 95B 25 mm) was used for imaging. Nerve endings in lung slices were acquired by confocal microscopy (Zeiss, LSM800), with a Z stack of the whole section (40 μm).

### Immunofluorescence analysis

Using ImageJ or Nikon Elements, circular ROI were manually defined around each neuron based on PGP9.5 fluorescence for JNC and SG slices. For confocal pictures of lung slices, an orthogonal projection of the z-stack was made, then a threshold-based method was used to define the area of nerve fibers and immune cells for each marker as well as their overlapping areas. Cultured cells were analyzed with a threshold-based method to define neurons based on tdTomato or PGP9.5 expression. Average fluorescence in ROI was exported and all further analysis were performed in Microsoft Excel. When spillover between colors occurred, a compensation was applied using single-stain controls.

### Neuron culture for RNA sequencing

Na_V_1.8^cre^::tdTomato^fl/wt^ mice were sacrificed and their JNC harvested. For each sample, JNC from an equal number of males and females were pooled and enzymatically dissociated, seeded (1×10^4^ / well) in 12 well plate (VWR, #10062-894), and cultured with IL-13 (100 ng/mL), IL-1β (100 ng/mL), TNF-α (100 ng/mL), BDNF (50 ng/mL) or vehicle. After 24 hours, the cells were then mechanically detached with a cell scraper and harvested in flow cytometry buffer (PBS, 2% FBS, 1 mM EDTA), filtered (70 μm), sorted by flow cytometry (FSC^hi^SSC^hi^tdTomato^hi^), and collected into Trizol.

### RNA sequencing

RNA libraries preparation and sequencing were carried out at the genomic platform of the Institut de Recherche en Cancérologie et en Immunologie (IRIC). Briefly, RNA quality was assessed using a Bioanalyzer (Agilent), and all preparations had an RIN>7.5. Libraries were prepared using the KAPA mRNA HyperPrep Kit (KapaBiosystems #KR1352). All barcoded samples were then sequenced with a Nextseq500 (Illumina) with 75-cycle single-end read.

Genome alignment and differential expression analysis were carried out (IRIC genomic platform). Sequences were trimmed for sequencing adapters and low quality 3’ bases using Trimmomatic version 0.35 and aligned to the reference mouse genome version GRCm38 (gene annotation from Gencode version M25, based on Ensembl 100) using STAR version 2.7.1. Gene expressions were obtained from STAR as readcounts and computed using RSEM to obtain normalized gene and transcript level expression in FPKM. Differential expression analysis for the various comparisons of interest were made using DESeq2 ^182^ and STAR readcounts. Further analysis and plots were made using RStudio or Microsoft Excel. Genes were considered as differentially expressed if their adjusted p-value (FDR) was less than 0.2.

### *In-silico* analysis of JNC neuron single-cell transcriptome

Prescott et al.^1^ generated single-cell sequencing data for jugular-nodose ganglia cells from 40 mice using the 10X Genomics platform. The data was downloaded from the NCBI Gene Expression Omnibus (GSE145216) and analyzed using Seurat. Neuronal cells were selected based on *Slc17a6* (VGLUT2) expression (raw count ≥ 2). A standard workflow was used for quality control, preprocessing, normalization, and clustering (resolution = 0.2, PCs = 1:30). *Phox2b* and *Prdm12* were used to identify nodose and jugular groups, while nociceptor neurons and low-threshold sensory neurons were defined based on their expression of *Scn10a* and *Scn1a* ^2^. Xhao et al.^7^ generated single-cell projection sequencing data for jugular-nodose ganglia cells from mice in which barcode expressing viruses were used to retrogradely label neurons innervating various internal organs. The data was downloaded from the NCBI Gene Expression Omnibus (GSE192987) and analyzed using Seurat. Neurons expressing at least one barcode (raw count ≥ 2) were selected. A standard workflow was used for quality control, preprocessing, normalization, and clustering (resolution = 0.75, PCs = 1:20).

### GSEA analysis

Gene set enrichment analysis ^183^ was performed to compare similarities between different sequencing results together using the GSEA software. To assess the preferential innervation of airways by scRNAseq neuron clusters, GSEA analysis was performed using the cluster markers identified in Prescott et al.’s dataset as genesets, and the DESeq2 counts of airway and visceral neurons as gene expression dataset. To compare AAI, BDNF, IL-1b, IL-13, TNF-a, and nerve injury sequencing data, gene signatures were constituted by selecting overexpressed genes (FDR<0.2) induced in each condition. For the nerve injury signature, RNAseq data from Cobos et al ^53^ was downloaded from NCBI Gene Expression Omnibus (GSE102937), reanalyzed by DESeq2 (**supplementary table 3**), and overexpressed genes (FDR<0.2) we selected. The enrichment of each gene signatures in each DESeq2 count datasets were measured by GSEA.

### Lung single-cell RNA-sequencing library preparation and sequencing

Mice lungs were harvested, minced with a razor blade, digested in Collagenase IV (1.6mg/ml) + DNAse I (10ug/ml) in DMEM buffer supplemented with Hepes (10mM) for 30 min. Lungs were triturated with a 1mL pipette and cells suspension were filtered (40um). 2 different mice were pooled for each sample. RBC were lysed (Gibco, #A10492-01) before staining with viability dye (Biolegend 423105) and anti-CD45 (103107). 500000 live immune cells were sorted on a FACS ARIA III (BD) and harvested in collection buffer (PBS+ 0.04% BSA + 50% FBS). Cells were then fixed, before single cell library preparation using Chromium 10x fixed RNA kits and instructions (#PN-1000414 and #PN-1000497). Libraries were subsequently sequenced on Illumina Novaseq X.

### Lung single-cell RNA-sequencing analysis

Reads were mapped to the mouse reference genome using Cell Ranger (Chromium) and analyzed using Seurat. Low RNA and dead cells were excluded (nFeature_RNA > 100 & percent.mt < 25). Total cells were clustered (dims = 1:20 and resolution = 0.5). 21 clusters were identified, and were assigned using common markers and the Tabula Muris reference. Markers used included: Cd19, S100a8, Cd3e, Klrk1, Cd200r3, Cd68, Itgax, Lgals3, H2-Ab1. Clusters were grouped as large cell types (B cells, T cells, NK cells, antigen presenting cells (APC), Granulocytes, Basophils, Alveolar macrophages, and endothelial cells) to compared expression between different conditions. To compare naïve and AAI conditions, a single library was prepared and the inflammation markers were defined using Seurat function FindMarkers. To compare gene expression between NPY1R conditional Knock-Out and littermate controls, 3 biological replicates per group were prepared. To maximise statistical robustness, a pseudobulk analysis was then used to compare groups. Gene expression was averaged for each cell type and for each sample, and Deseq2 was used to identify differentially expressed genes, while GSEA was used to define gene ontology gene sets. Since a very low number of Basophils, Alveolar macrophages, and endothelial cells were sequenced, those were not used for the pseudobulk analysis. To focus the analysis on T cells specifically, T cells were subsetted, and Seurat functions were used to define the different groups of t cells (dims =1:40, resolution =, 1). Since V(D)J genes were driving excessive clustering of the T cells, those were not taken into account for the clustering. T cell clusters were defined based on various common markers including: *Cd3e, Cd4, Cd8, Cd8b1, Il1rl1, Il13, Il17a, Trdc, Rag1, and Klrc1* (**SF. 6C**).

## Data availability

Bulk and single-cell RNA-sequencing raw and processed data have been deposited in the NCBI’s gene expression omnibus (GSE223355; *single cell number pending*). Processed data can also be accessed in the **supplementary tables 1–10**. Additional information and raw data are available from the lead contact upon reasonable request.

## Statistics

No data were excluded. P *values ≤ 0.05* were considered statistically significant. One-way ANOVA, two-way ANOVA, and Student t-tests were performed using GraphPad Prism. DESeq2 and Seurat analysis and statistics were performed using RStudio.

## Replicates

Replicates (*n*) are described in the figure legends and represent the number of animals for *in vivo* data. For *in vitro* data, replicates can either be culture wells or dishes, animals, fields-of-view (microscopy), or neurons (patch-clamp), but always include different preparations from different animals to ensure biological reproducibility.

## Acknowledgements

ST work is supported by the Canada Research Chair program (950-231859). Canadian Institutes of Health Research (CIHR; 407016, 461274, 461275), Canadian Foundation for Innovation (44135), Canadian Cancer Society Emerging Scholar Research Grant (708096), Knut and Alice Wallenberg Foundation (KAW 2021.0141, KAW 2022.0327), Swedish Research Council (2022-01661), Natural Sciences and Engineering Research Council of Canada (NSERC; RGPIN-2019-06824), and NIH/NIDCR (R01DE032712). RB’s work is supported by CIHR (169160) and NSERC (RGPIN-2017-06871). SB received a fellowship from the centre interdisciplinaire de recherche sur le cerveau et apprentissage (CIRCA). We thank the genomic platform of the Institut de recherche en Cancérologie and Immunologie (IRIC), P. Gendron for the bioinformatic analysis, and R. Lambert and J. Huber for library preparation and sequencing. We also thank P.D.B. Rosa for graphic design.

