## Supplementary figures and images for "Cytokines reprogram airway sensory neurons in asthma"

### Supplemental Fig. 1

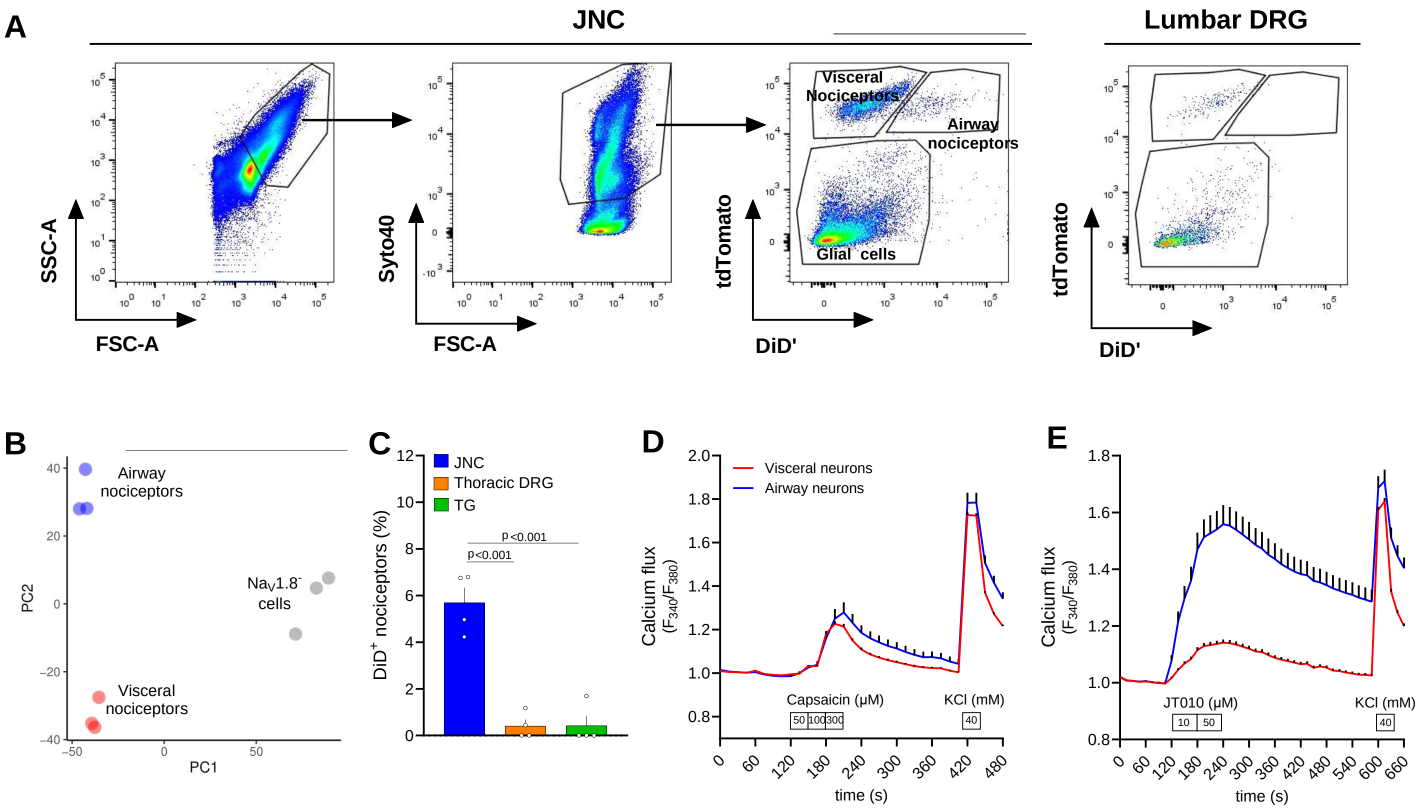

### Supplemental Fig. 2

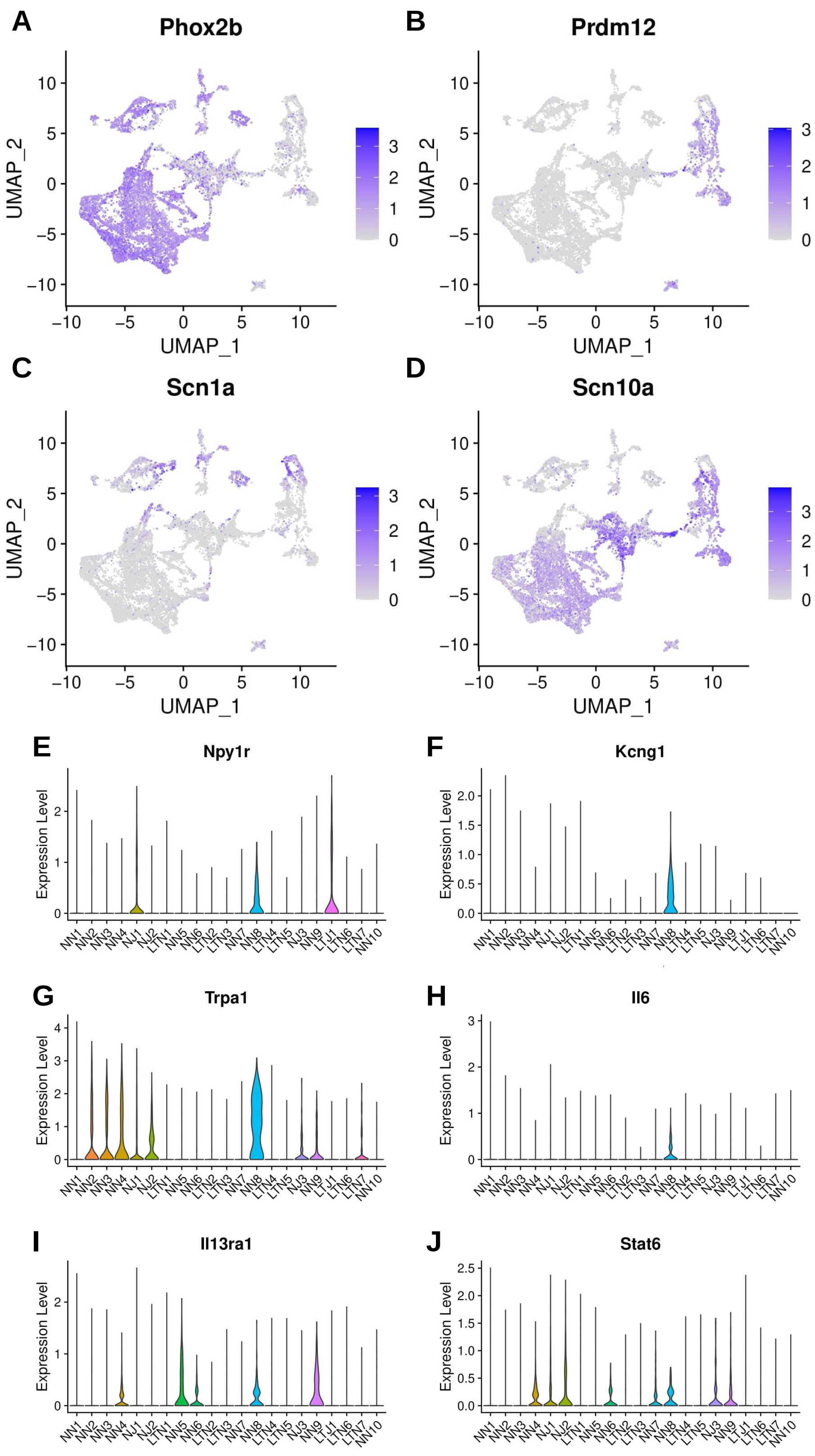

### Supplemental Fig. 3

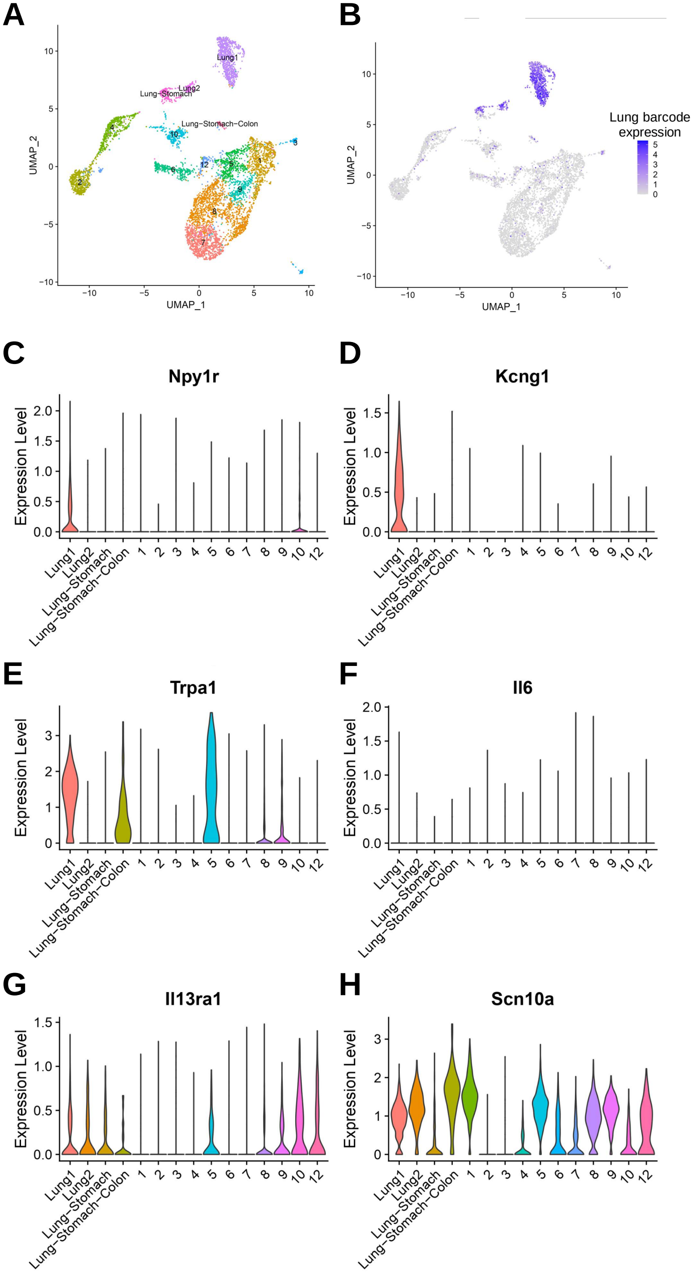

### Supplemental Fig. 4

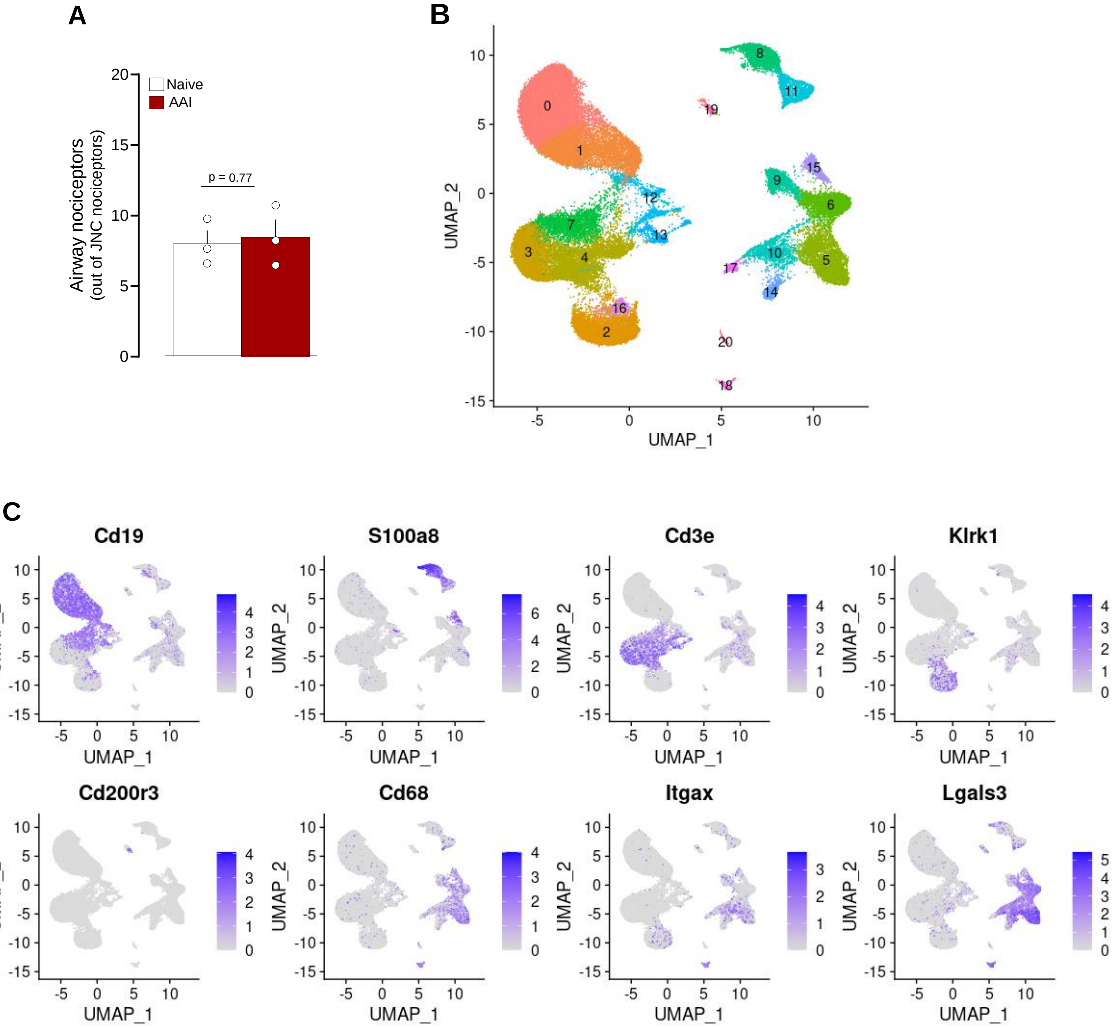

### Supplemental Fig. 5

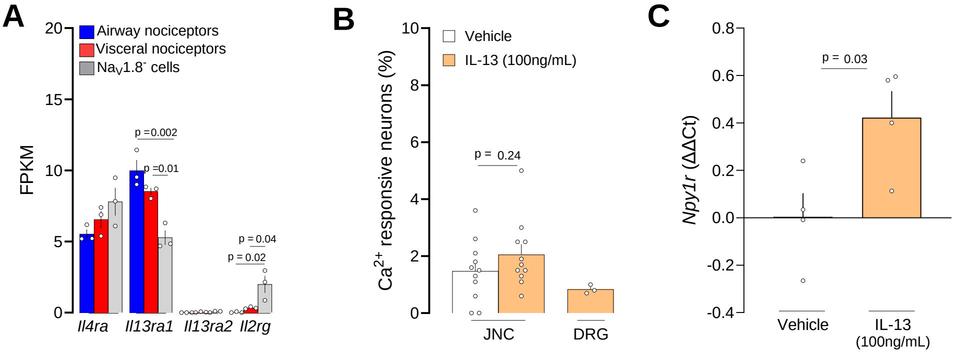

### Supplemental Fig. 6

A

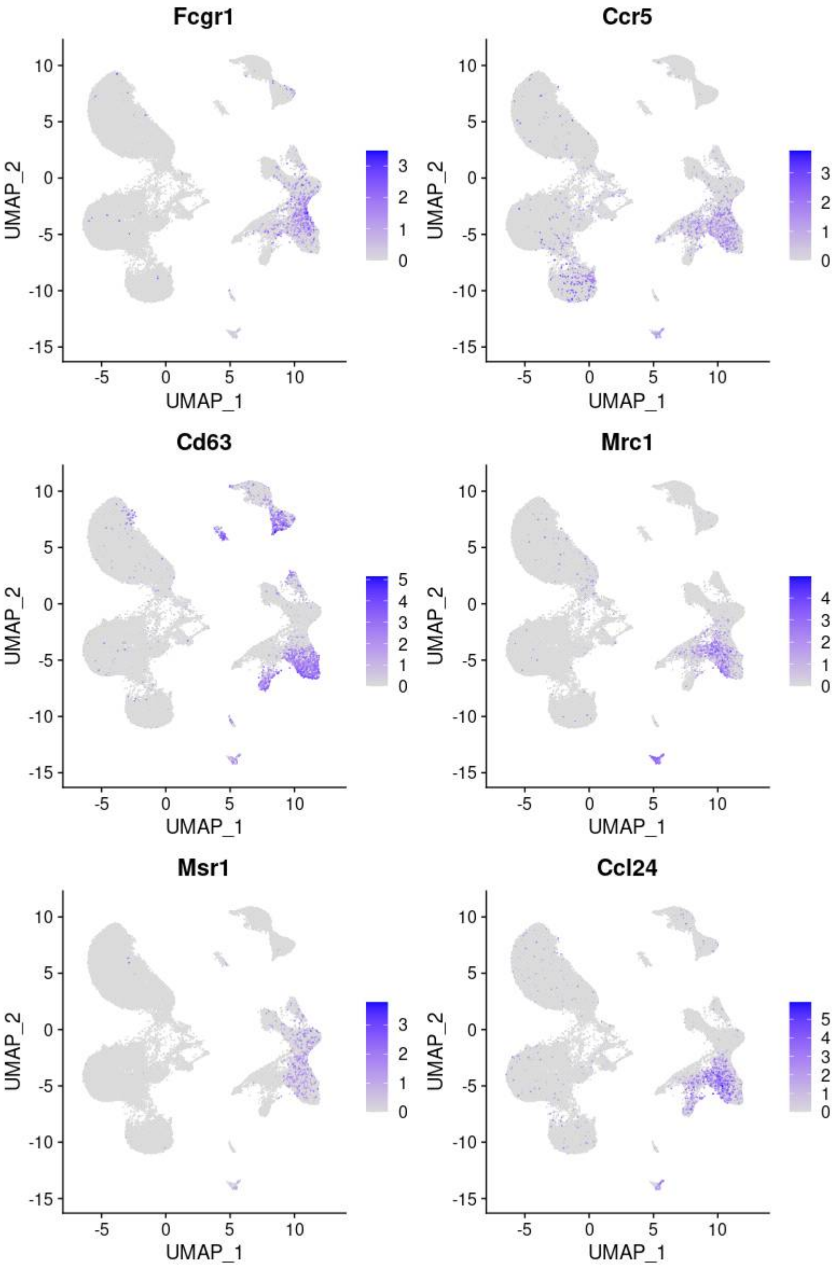

### Supplemental Fig. 7

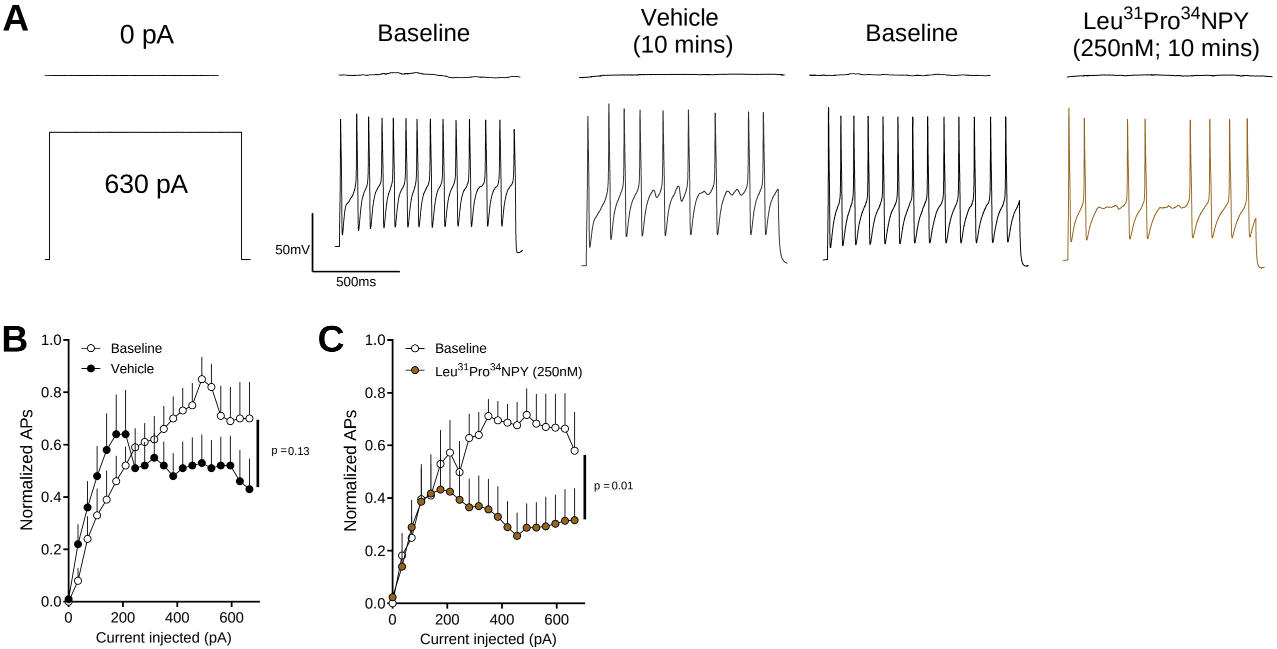

### Supplemental Fig. 8

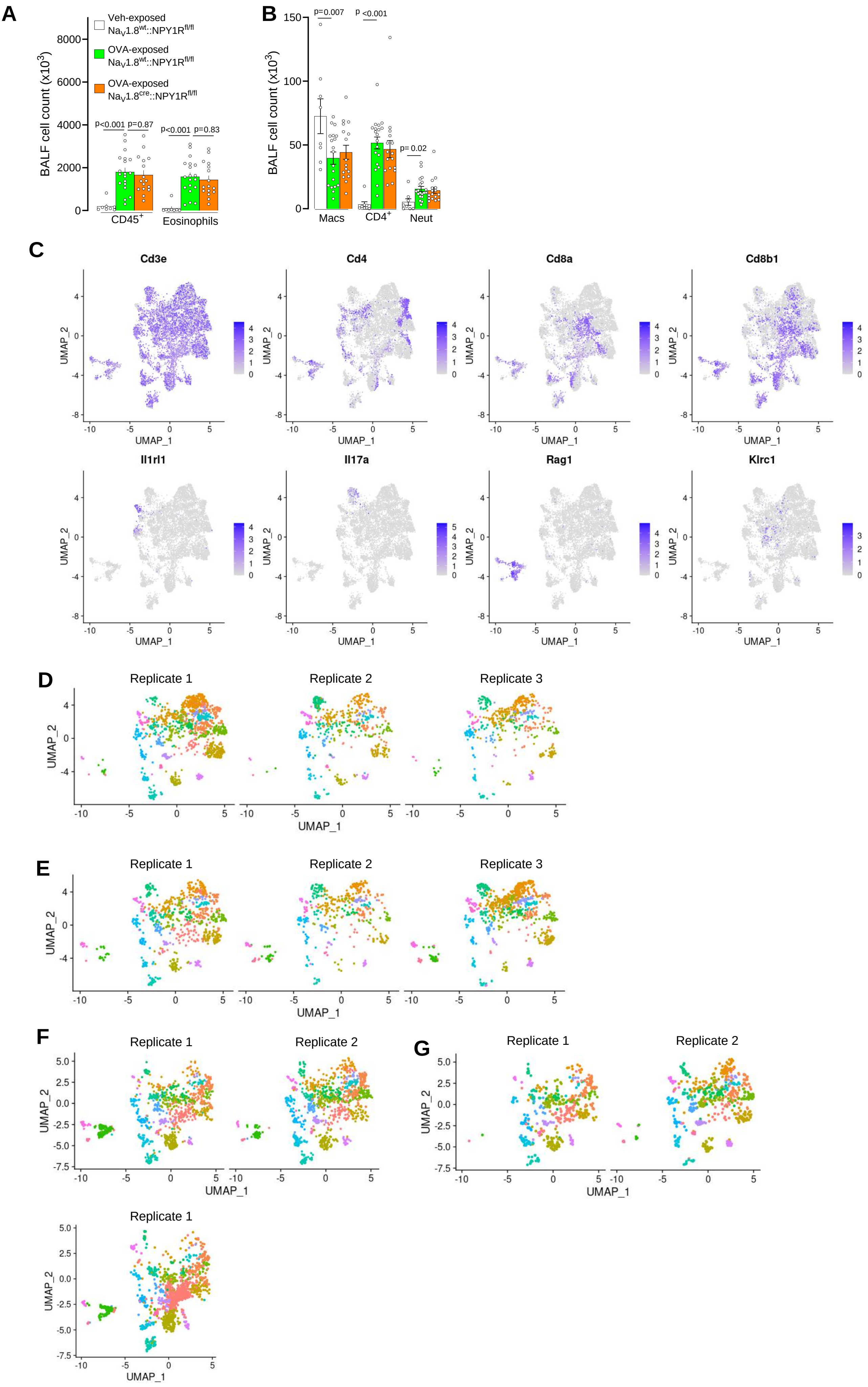
